# Single Cell RNAseq Reveals A Critical Role of Chloride Channels in Airway Development

**DOI:** 10.1101/735969

**Authors:** Mu He, Bing Wu, Daniel D. Le, Wenlei Ye, Adriane W. Sinclair, Valeria Padovano, Yuzhang Chen, Kexin Li, Rene Sit, Michelle Tan, Michael J. Caplan, Norma Neff, Yuh Nung Jan, Spyros Darmanis, Lily Y. Jan

**Author notes:** these authors contributed equally. Correspondence (M.H), (S.D), (L.Y.J).

## Abstract

The conducting airway forms a protective mucosal barrier and is the primary target of airway disorders. To better understand how airway developmental programs are established to support air breathing and barrier functions, we constructed a single-cell atlas of the human and mouse developing trachea. In this study, we uncover hitherto unrecognized heterogeneity of cell states with distinct differentiation programs and immune features of the developing airway. In addition, we find ubiquitous expression of *CFTR* and *ANO1/TMEM16A* chloride channels in the embryonic airway epithelium. We show that genetic inactivation of TMEM16A leads to airway defects commonly seen in cystic fibrosis patients with deficient CFTR, alters the differentiation trajectory of airway basal progenitors, and results in mucus cell hyperplasia and aberrant epithelial antimicrobial expression. Together, our study illuminates conserved developmental features of the mammalian airway and implicates chloride homeostasis as a key player in regulating mucosal barrier formation and function relevant to early onset airway diseases.

## INTRODUCTION

The highly conserved respiratory system of air breathing animals represents a major interface between internal organs and the outer environment. In the course of a typical human lifespan, approximately 200 to 400 million liters of air are conducted via the respiratory system (Ganesan et al., 2013; Rackley and Stripp, 2012). While airway function has been adapted for organismal physiology and aging (Sharma and Goodwin, 2006), it remains vulnerable to deleterious genetic and environmental factors. Chronic pulmonary conditions, such as asthma, chronic obstructive pulmonary disease (COPD), and cystic fibrosis (CF), primarily target the trachea and bronchi.

These disorders are characterized by mucus obstruction and repetitive infections and inflammation, often leading to severe airway remodeling and respiratory failure (Bergeron et al., 2010; James and Wenzel, 2007). While the pathogenesis of many of these chronic conditions remains unclear, early manifestation of airway symptoms in young patients with cystic fibrosis and asthma suggests that alterations of airway development can have a profound impact on the respiratory function later in life (Holt and Sly, 2012; Larson and Cohen, 2005; Taylor et al., 2019).

The architecture of the trachea and bronchi is largely conserved in mammals (Hogan et al., 2014; Rackley and Stripp, 2012). The airway epithelial tube is surrounded by rigid cartilage rings on the ventral side and elastic smooth muscles on the dorsal side. Luminal cells, such as multiciliated cells and secretory cells, are differentiated from basal progenitor cells during embryogenesis (Rock et al., 2010). After animals start breathing, these luminal cells of the conducting airway form a protective mucosal barrier, clear inhaled pathogens, and maintain airway fluid balance, whereas basal progenitors support tissue homeostasis and regeneration in response to injury (Hogan et al., 2014; Rackley and Stripp, 2012). Building on decades of genetic and developmental studies, recent transcriptome analyses utilizing single cell RNA sequencing (scRNAseq) have systematically characterized airway cell types and lineage networks within the adult airway under normal and chronic inflammatory conditions (Lee et al., 2017; Montoro et al., 2018; Ordovas-Montanes et al., 2018; Plasschaert et al., 2018; Vieira Braga et al., 2019). Whether the same programs for adult airway regeneration are also operating during embryogenesis remains an open question.

To define the cellular processes important for airway development and disease manifestations that begin early in life, we used scRNAseq to profile mouse embryonic and neonatal trachea as well as human fetal trachea. We uncovered conserved cell types implicated in monogenic and complex-trait airway diseases and defined cell states associated with epithelial cell differentiation. In addition, we identified a novel cilia-secretory hybrid state, which exhibit features of ciliated cells and secretory cells and are predominantly present in the neonatal trachea.

To probe the cellular origin of mucous hyperplasia, a hallmark of many pulmonary conditions, we analyzed the developmental landscape of the mouse trachea in the absence of *Ano1/Tmem16a*, a gene encoding for a calcium-activated chloride channel with diverse physiological functions, including neuromodulation in mammals and the inhibition of polyspermy in amphibian eggs (Caputo et al., 2008; Schroeder et al., 2008; Wozniak et al., 2018; Yang and Jan, 2017; Yang et al., 2008). *Tmem16a* is expressed in the mouse airway, and *Tmem16a* KO mice exhibit tracheomalacia and accumulation of mucus in the trachea (Rock et al., 2008; 2009). TMEM16A-mediated chloride conduction is hypothesized to serve as an alternative chloride secretion pathway in the mouse trachea, and may account for the lack of CF-like phenotypes in the airway of *Cftr* knockout mice, in which CFTR channel activity is abolished (Clarke et al., 1994; McCarron et al., 2018; Sondo et al., 2014). Here we show that *Cftr* and *Tmem16a* are widely expressed in epithelial cells during embryogenesis of mice and humans. We also showed that *Tmem16a* in mice plays a key role in controlling the differentiation of basal progenitors into the secretory lineage. Lack of *Tmem16a* results in abnormal accumulation of intermediate secretory cells, which may account for the mucosal hyperplasia seen in the *Tmem16a^-/-^* mutants. In addition, *Tmem16a* mutants display abnormal immune profiles including increased expression of pro-inflammatory cytokines and decreased expression of antimicrobial molecules. In summary, our study establishes a cell atlas for the developing airway of mice and humans, revealing cell states and associated genes that have not been previously described. We also present a tractable mouse model for understanding the cellular processes controlled by chloride homeostasis in the formation of mucosal barrier relevant to early onset airway diseases.

## RESULTS

### Cellular Composition and Molecular Signatures of the Developing Mouse Trachea

We performed scRNAseq of tracheal tissue collected at embryonic days 15 and 16 (E15 and E16), during which airway luminal cells begin to appear, as well as at postnatal days 1 and 4 (P1 and P4), when airflow increases as the animals begin to breathe and the mucosal barrier function of the airway develops (Figure 1A) (Ganesan et al., 2013; Swarr and Morrisey, 2015). A total of approximately 16000 cells that are wild-type for the *Ano1* gene locus passed quality control (Figure S1A), and were subsequently subjected to dimensionality reduction by principal component analysis (PCA) using the top 1738 overdispersed genes, followed by nearest-neighbor graph-based clustering on the PC space (Butler et al., 2018). We assigned a cellular identity to each cluster using a collection of known marker genes for mouse airway cell types that was cross referenced with differentially expressed genes between clusters (Figures 1B-C and S1B). For E15 and P4 samples, we performed a lineage tracing experiment by using *Shh^Cre^/R26^mT/mG^* mice in order to distinguish airway epithelial cells derived from the foregut endoderm from cells of other lineages (Swarr and Morrisey, 2015). In agreement with the established lineage relationships of the airway endoderm, GFP^+^ cells derived from *Shh*^+^ endoderm consistently expressed the epithelial marker *Epcam*. These *Epcam^+^* cells were annotated as 1) basal cells, 2) luminal cells including ciliated cells and secretory cells, and 3) a newly identified cilia-secretory hybrid state, which simultaneously expressed genes characteristic of both ciliated cells and secretory cells (Figures 1C and S1C). RFP^+^ cells derived from non-*Shh* expressing lineages consisted of a large collection of mesenchymal cells, muscle cells, endothelial cells, immune cells, Schwann cells, and neuronal cells (Figures 1C and S1C).

**Figure 1.**
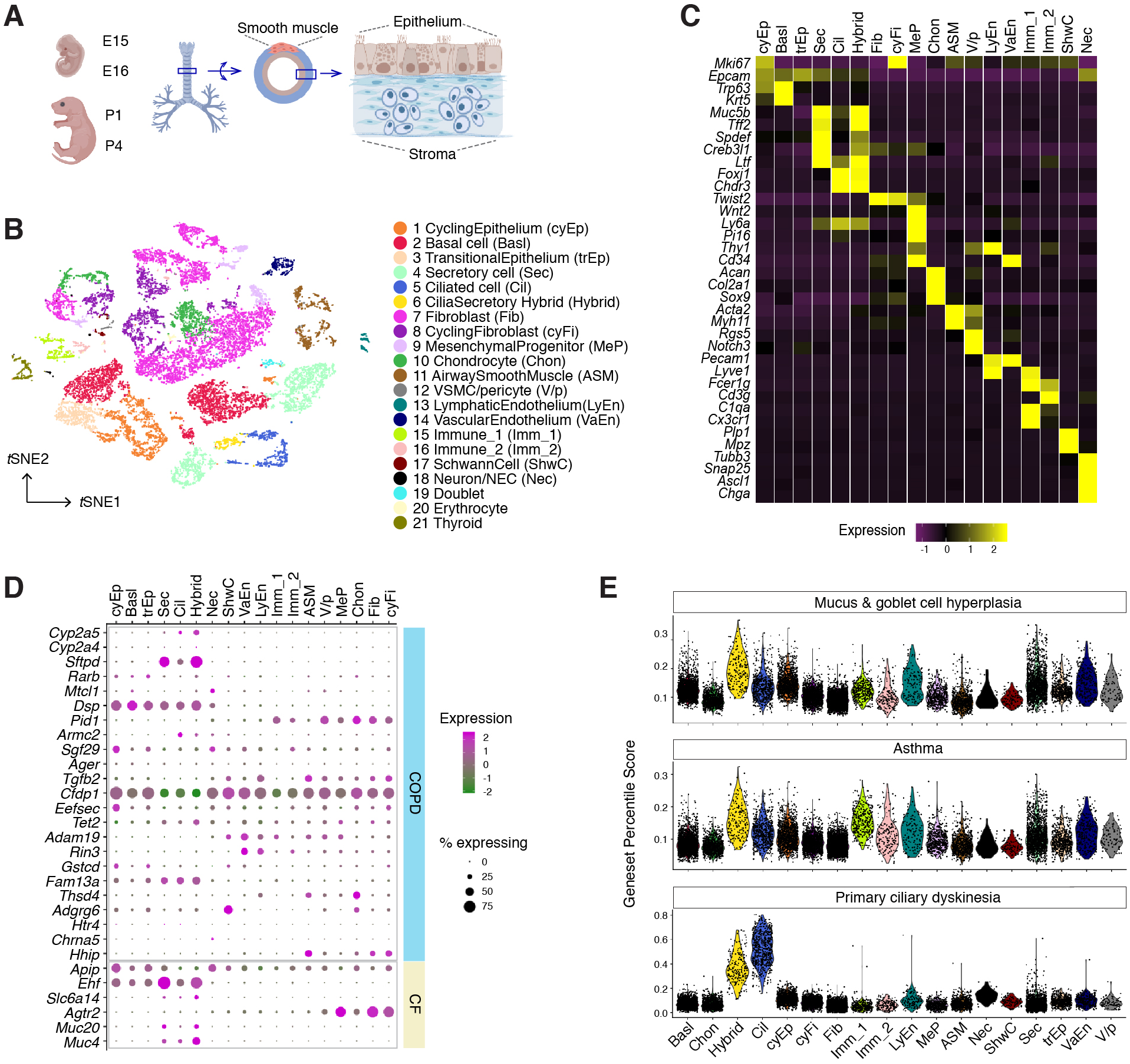
Single Cell Atlas of the Developing Mouse Trachea. (A) Cartoons of the mouse trachea anatomy. Four developmental stages are included in this study: embryonic day 15 (E15), E16, postnatal day 1 (P1), and P4. (B) Tracheal cells from wild-type mice visualized on a *t*-distributed stochastic neighbor embedding (*t*SNE) plot. Colors indicate cell types and states identified in this study. (C) Heat map illustrating the average expression of marker genes for each cluster. Gene expression has been normalized, log-transformed, and z-score transformed. Gene markers for erythrocytes and thyroid cells are included in Figure S1E and S1F. (D) Dot plot depicting expression patterns of genes implicated in the disease severity of COPD and CF. The size of the dot encodes the percentage of cells expressing the gene, while the color encodes the mean of expression level which has been normalized, log-transformed, and z-score transformed. (E) Single cell geneset percentile score (scGPS) of disease-associated genes curated from the Online Mendelian Inheritance in Man (OMIM) database. Each dot represents a cell. Colors indicate cell clusters. Full lists of genes used are in Table S1.

We also found a small cluster of cells with consistently high doublet scores (Wolock et al., 2019) that simultaneously expressed both *Epcam* and the mesenchymal marker *Twist2* (Figure S1D). In addition, we identified two non-tracheal cell clusters, including a thyroid cluster marked by Thyroglobulin (*Tg*) and *Pax8* (Figure S1E), and an erythrocyte cluster marked by *Alas2* and *Hba-a2* (Figure S1F). Doublets, thyroid cells, and erythrocytes were excluded from subsequent analyses.

### Molecular Taxonomy of Mesenchymal Cells of the Developing Mouse Trachea

Our dataset comprises a collection of diverse mesenchymal cell types, many of which have not been characterized at single cell resolution. Vascular smooth muscle cells and pericytes are identified based on the expression of *Notch3* and *Rgs5*, whereas airway smooth muscle cells express higher levels of *Myh11* and *Acta2* (Figure 1C). Endothelial cells express *Pecam1* and can be further grouped into lymphatic endothelial cells based on the expression of *Lyve1* and *Thy1*, and vascular endothelial cells based on the expression of *Cd34*. We identified two immune cell clusters, including a population of *Fcerig*^+^/*Cd3g*^+^ T cells and a population of *Cx3cr1*^+^/*C1qa*^+^ monocytes (Figure 1C). Our dataset includes a cluster of *Wnt2*^+^ mesenchymal cells which persist across all time points included in this study. These cells are marked by *Pi16*, *Cd34*, and *Ly6a* (*Sca-1*) (Figure 1C), similar to the molecular signatures of adipose progenitor-like cells (Baryawno et al., 2019; Sanchez-Gurmaches et al., 2016). Because *Wnt2*^+^ lineages can serve as cardiopulmonary progenitors and define a mesenchymal alveolar niche important for self-renew and repair in the lung (Peng et al., 2013; Zepp et al., 2017), we annotated this *Wnt2^+^*/*Cd34^+^*/*Ly6a^+^* cluster as mesenchymal progenitors, which may generate the reservoirs of mesenchymal cell types during development and tissue repair.

We also captured a cluster marked by *Mpz* and *Plp1*, which are expressed in Schwann cells derived from the neural crest (Dyachuk et al., 2014; Espinosa-Medina et al., 2014; Feltri et al., 1999), as well as a small cluster distinguished by *Snap25*, *Ascl1*, *Chga*, which are molecular signatures for neuroendocrine cells (Nec) present in the adult mouse trachea (Montoro et al., 2018; Plasschaert et al., 2018) (Figures 1C and S1G). These Nec express transcription factors *Phox2a* and *Phox2b* required for cell fate specification in the parasympathetic ganglia (Figure S1G) (Dyachuk et al., 2014; Espinosa-Medina et al., 2014), indicating that trachea Nec may originate from a lineage different from the pulmonary Nec found in the vertebrate lung (Hockman et al., 2017).

### Mapping Monogenic and Complex Trait Disease-associated Genes to Cell Types of the Airway

Next we evaluated putative cell types from which complex airway may disorders arise. We examined the expression landscape of genetic risk loci associated with COPD and pulmonary fibrosis (Sakornsakolpat et al., 2019), as well as modifier loci associated with cystic fibrosis, a monogenic disease caused by mutations in the *CFTR* gene (Corvol et al., 2015). Many of the airway disease associated genes are expressed by cells of the airway epithelium, including modifier loci for CF severity, such as *Muc20* and *Ehf*, as well as risk genes shared between COPD and pulmonary fibrosis, such as *Fam13a* and *Dsp*. Nonetheless, expression of disease associated genes is not restricted to the epithelium (Figure 1D). For example, *Agtr2*, a CF modifier involved in various pulmonary functions, showed abundant expression in airway fibroblasts. COPD loci *Rin3* and *Adam19* are enriched in endothelial cells, whereas *Tgfb2*, *Thsd4*, and *Hhip* are enriched in smooth muscle cells (Figure 1D). Overall, risk genes implicated in these airway diseases are expressed in various cell types, and our dataset enables mapping of each disease mediator to its contributing cellular source.

Using scGPS (single-cell Geneset Percentile Score), an expression enrichment analysis that utilizes sets of genes underlying certain biological and pathological processes to infer functional profiles for each cell, we assessed whether the function of specific cell types can be inferred from the expression patterns of genes whose loss of function results in airway pathophysiology. Using genes curated by OMIM, we first assessed enrichment patterns for genes implicated in mucus and goblet cell hyperplasia (Figure 1E; Table S1), which are typically present in inflammation and infection and are classic symptoms for COPD and CF. Scores for mucosal hyperplasia are higher in cilia-secretory hybrid cluster, and are also elevated in T cells, endothelial cells, and vascular smooth muscles (Figure 1E). Similar enrichment profiles are observed for asthma associated genes, indicating shared cellular drivers for goblet cell hyperplasia and asthma (Figure 1E). Primary ciliary dyskinesia (PCD) is a monogenic disorder associated with impaired motile cilia function. Patients of PCD show persistent rhinitis and chronic respiratory tract infections, but are often initially diagnosed as asthma or bronchiectasis (Lucas et al., 2014; Shapiro et al., 2018). The PCD scores are higher overall for ciliated cells and cilia-secretory hybrid cells, confirming that ciliated cells are primary targets for PCD. The results of our scGPS analysis provide functional validation of our cell type annotation, and help disambiguate cell-type involvement in the manifestation of monogenic and complex-trait diseases. Interestingly, the cilia-secretory hybrid cells show prominent enrichment scores in multiple airway conditions (Figure 1E), indicating a critical role of this novel cell state in airway function and pathogenesis.

### The Developmental Landscape of the Mouse Trachea from Embryogenesis to Neonates

The temporal aspects of our developing mouse trachea atlas allowed the identification of cell types and molecular programs required for the morphogenesis and physiological maturation of the conducting airway. Because epithelial cells of the mucosal barrier are implicated in the manifestation of early onset airway diseases, we examined the developmental landscape of epithelial cell types during normal tracheal development.

First we identified subpopulations of cells within each epithelial cell type, and constructed gene modules characteristic for each subpopulation based on differential expression analysis. We used this information to annotate differentiation states that correlate closely with distinct developmental stages (Figure 2A). Our data show that coordinated differentiation programs of luminal cell types (ciliated and secretory) were initiated between E15 and E16. *Spdef* and *Creb3l1*, two transcription factors that specify secretory cells (Chen et al., 2009; Fox et al., 2010; Gregorieff et al., 2009) and *Foxj1* which specifies ciliated cells (You et al., 2004; Yu et al., 2008), were upregulated at E16 compared to E15 (Figure 2B). Conversely, the number of cycling cells, reflected by a calculated cell cycle score, was diminished as differentiated cells began to emerge (Figure 2C). At E15, the broad presence of *Trp63* but low level of *Krt5*, and marked expression of *Id2*, *Id3*, *Wnt7b*, and *Cldn6* indicate that undifferentiated cells dominate the *Epcam*^+^ epithelium (Figure 2D i). Id proteins are shown to promote proliferation and to inhibit differentiation, and their gene expression is reduced upon lineage commitment (Becker-Herman et al., 2002; Kowanetz et al., 2004; Roschger and Cabrele, 2017). *Wnt7b* is a Wnt ligand expressed in airway epithelium and regulates lung mesenchymal and vascular growth (Shu et al., 2002; Wang et al., 2005), and is implicated in the differentiation of hair follicle stem cells and pancreatic progenitors (Afelik et al., 2015; Kandyba and Kobielak, 2014). *Cldn6* is a tight junction protein of which the expression peaks prior to the differentiation of airway epithelium (Jimenez et al., 2014; Sugimoto et al., 2013). When luminal cells begin to emerge at E16, *Trp63*^+^ basal cells switch to a different expression program, consisting of *Krt5*, *Krt15* (Moll et al., 2008), *Aqp3,* and *Aqp4*, which may play important roles in the epithelial barrier function (Kreda et al., 2001; Matsuzaki et al., 2009) (Figure 2D i).

**Figure 2.**
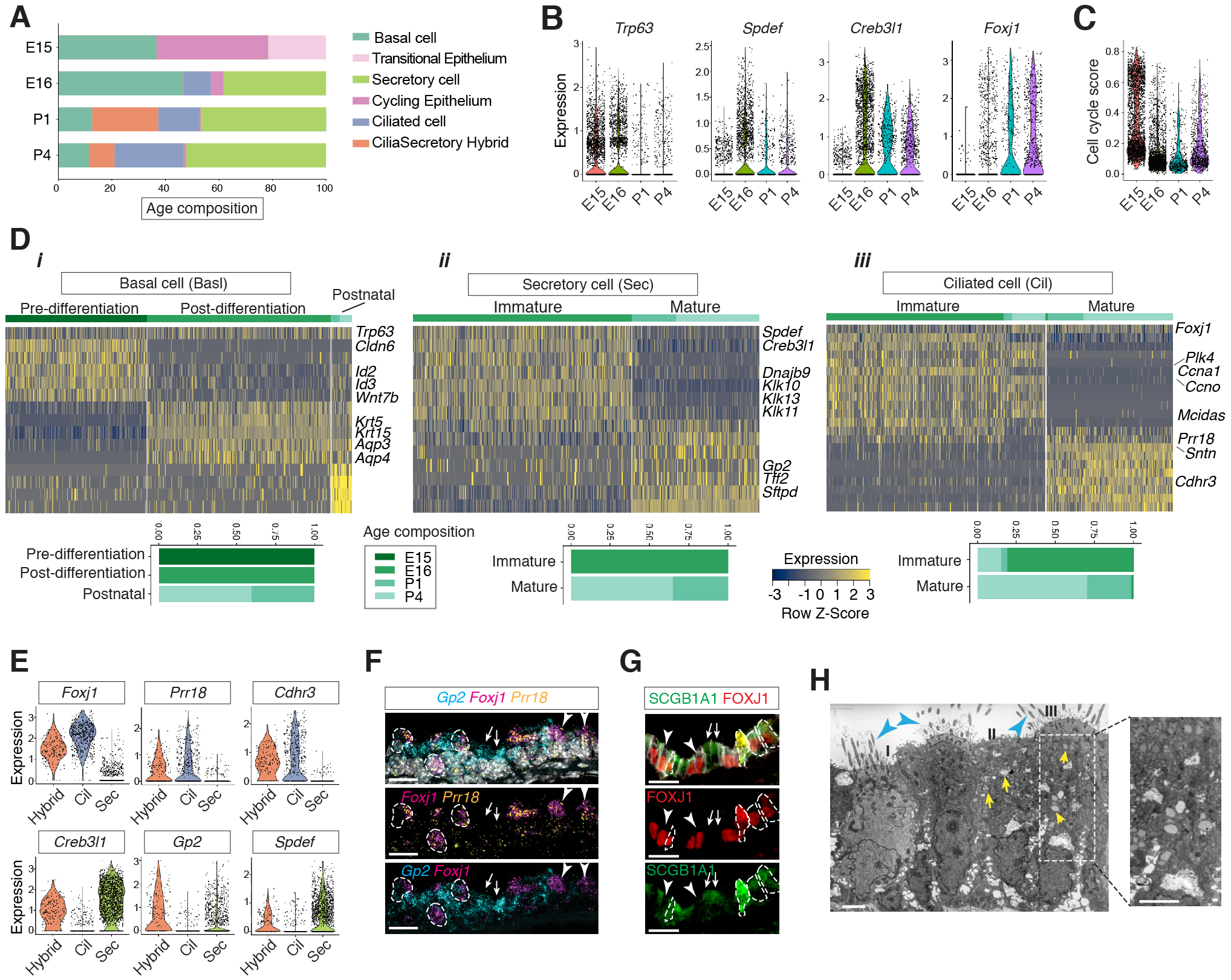
Developmental Landscape of Tracheal Epithelial Cells. (A) Horizontal bar graphs showing the cellular composition of tracheal epithelial cells from different time points. (B) Violin plots showing the expression of *Trp63*, *Spdef*, *Creb3l1*, and *Foxj1* in epithelial cells at different developmental time points. Gene expression has been normalized and log-transformed. (C) scGPS showing cell-cycle gene expression in epithelial cells at different time points. The cell cycle gene list can be found in Table S2. (D) Heat maps showing transcriptional profiles of tracheal basal, ciliated, and secretory cells across different developmental time points, with age compositions of corresponding cell states showed in horizontal bar plots. Gene expression has been normalized, log-transformed, and z-score transformed. A complete genelist for the heat maps can be found in Table S2. (E) Violin plots showing the expression levels of *Foxj1*, Prr18, *Cdhr3*, *Creb3l1*, *Spdef*, and *Gp2* in ciliated cells, secretory cells, and cilia-secretory hybrids. Gene expression has been normalized and log-transformed. (F) Fluorescent *in situ* hybridization (FISH) of *Foxj1* (magenta), *Gp2* (blue), and *Prr18* (yellow) mRNA in P3 wild-type trachea. Hybrid cells expressing all three markers are indicated by dashed circles. Foxj1^+^ Prr18^+^ ciliated cells are indicated by arrowheads. Gp2^+^ secretory cells are indicated by arrows. Nuclei are marked by DAPI (grey). Scale bar indicates 20 μm. (G) Fluorescent immunostaining of FOXJ1 (red) and SCGB1A1 (red) in P3 wild-type trachea. Hybrid cells expressing both markers are indicated by dashed circles. FOXJ1^+^ ciliated cells are indicated by arrowheads. SCGB1A1^+^ secretory cells are indicated by arrows. Cell membranes are marked by E-cadherin (grey). Scale bar indicates 20 μm. (H) Transmission electron microscopy (TEM) images of P0 tracheal epithelial cells. I: ciliated cell; II: secretory cell; III; cilia-secretory hybrid. Intracellular vesicles are indicated by arrows in yellow. Motile cilia are indicated by arrowheads in blue. Scale bar indicates 2 μm.

At E16, in addition to *Spdef* and *Creb3l1*, the secretory program showed a pronounced expression of *Cited1*, *Dnajb9*, as well as Kallikrein peptidases, including *Klk10*, *Klk11*, and *Klk13* (Figure 2D ii), all of which have not been previously linked to the airway secretory program. Conversely, the neonatal secretory program was distinguished by *Gp2* and *Tff2*, which are important mucosal proteins and markers for goblets cells found in the adult airway (Hase et al., 2009; Montoro et al., 2018; Nikolaidis et al., 2006; Wills-Karp et al., 2012), as well as *Sftpd*, which encodes a surfactant protein and is involved in the innate immune program (Brandt et al., 2008; Mackay et al., 2016) (Figure 2D ii).

Ciliated cells, which form motile cilia, are essential for mucus clearance. In our dataset, both embryonic and postnatal ciliated cells expressed *Foxj1* (Figure 2D iii). Immature ciliated cells, which are predominantly present during embryogenesis, were characterized by *Ccna1*, *Ccno*, *Mcidas*, and *Plk4* (Figure 2D iii). These genes are cell cycle regulators operating in a hierarchical order upstream of *Foxj1* to promote a ciliated cell fate (Kyrousi et al., 2015; Spassky and Meunier, 2017; Vladar et al., 2018). Human mutations in *CCNO* and *MCIDAS* impair cilia formation and result in congenital mucociliary clearance disorder (Boon et al., 2014; Kyrousi et al., 2015; Wallmeier et al., 2014). In contrast, postnatal mature ciliated cells upregulate *Sntn*, which encodes a phosphatidylserine binding protein localizing to the tip of motile cilia (Kubo et al., 2008), and express various membrane receptors, such as *Cdhr3* and *Ldlrad1*, which are involved in rhinovirus infection (Basnet et al., 2019) (Figure 2D iii). An enrichment of these molecular markers thus reflects a structural and functional maturation of postnatal ciliated cells.

We did not observe Tuft cells in our dataset. Using DCLK1, a microtubule associated kinase that labels Tuft cells in various tissues including the trachea (Gerbe et al., 2009; Ting and Moltke, 2019), we found that DCLK1^+^ Tuft cells emerged postnatally and were present sparsely in the newborn trachea (Figures S2A and S2B).

### Identification of a Novel Cilia-Secretory Hybrid Cell State

Among luminal cell types, we identified a cell cluster exhibiting two sets of gene modules, the *Foxj1/Cdhr3*-associated cilia module and the *Gp2*/*Creb3l1*-associated secretory module (Figures 2E and S2C). This novel cell state was mostly found in our postnatal dataset (Figures 2A and S2D), and exhibited elevated expression of genes implicated in mucus and goblet cell hyperplasia as the animals develop (Figure S2E). While absent from normal adult airway of mice and humans, a cilia-secretory hybrid state has been observed in human samples with allergic conditions (Ordovas-Montanes et al., 2018; Vieira Braga et al., 2019), indicating that such hybrid state may appear under physiological conditions that favors inflammation, including the Th2 dominant immunity found in newborns (Iwasaki and Medzhitov, 2015; Kollmann et al., 2017; Levy, 2007; Simon et al., 2015).

Using neonatal tracheal samples, we validated the presence of a cilia-secretory hybrid cell type by RNA FISH (f*luorescent in-situ hybridization)* analysis of *Foxj1*, *Gp2* and *Prr18*, a novel ciliated cell marker (Figures 2D, 2E, and 2F). Based on immunofluorescence staining, these hybrid cells expressed FOXJ1 and a secretory cell marker SCGB1A1 (Figure 2G) (Rawlins et al., 2009; Zhang et al., 1997). Transmission electron microscopy (TEM) revealed that a subset of luminal cells exhibit both characteristic cilia axoneme and intracellular vesicles, indicating that hybrid cells indeed possess two sets of machineries required for motility and secretion, respectively (Figure 2H). To probe the developmental origin of these hybrid cells, we assessed their presence in a mouse mutant lacking *Pofut1*, which encodes an enzyme required for Notch ligand processing (Stahl et al., 2008). Because the Notch pathway is essential for cell fate decision in the airway, *Pofut1* mutants fail to produce secretory cells and contain predominantly ciliated cells in the trachea (Tsao et al., 2009) (Figure S2F). Compared to the wild-type, most, if not all, *Pofut1* mutant epithelial cells were labeled by acetylated-α-tubulin that marks motile cilia axoneme and expressed *Foxj1,* while *Gp2* was present in a fraction of cells that also expressed *Foxj1*, indicating that hybrid cells most likely arise from the ciliated cell lineage (Figure S2F).

### Inactivation of *Tmem16a* Leads to Mucus Cell Hyperplasia

In contrast to the adult trachea, where *Cftr* expression is restricted to rare ionocyte cells (Montoro et al., 2018; Plasschaert et al., 2018; Vieira Braga et al., 2019), we observed that *Cftr* expression was abundant in the undifferentiated mouse tracheal epithelium at E15 (Figures 3A and 3B). As luminal cell types started to emerge, *Cftr* expression decreased and became more restricted to secretory cells. At P4, *Cftr* was only detected at low levels and in a limited number of cells, while the expression of *Ano1*/*Tmem16a* persisted (Figures 3B and S3A).

**Figure 3.**
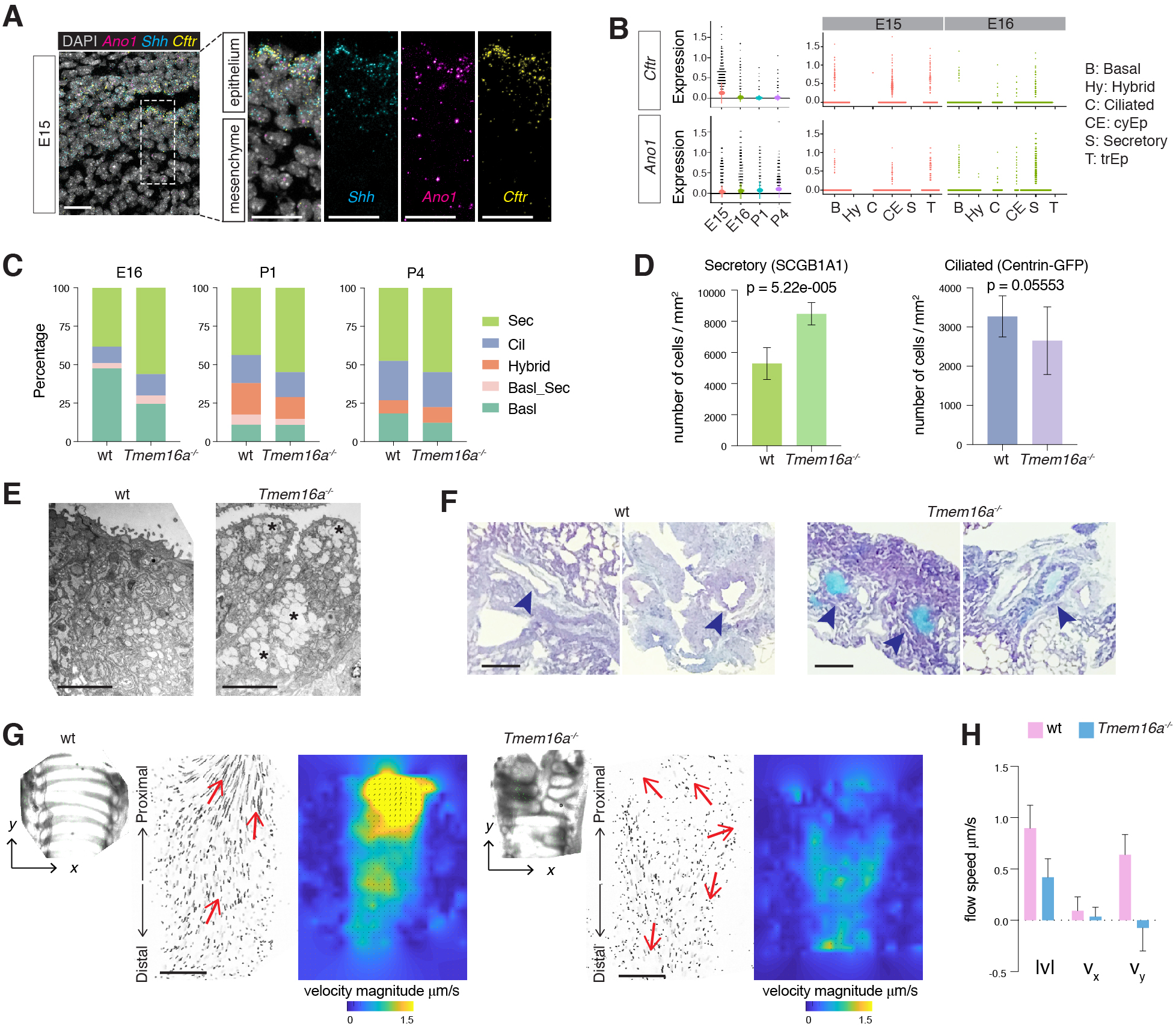
Airway Defects in *Tmem16a^-/-^* KO Mutants Associated with Secretory Cell Hyperplasia. (A) Expression of *Shh*, *Cftr*, and *Ano1/Tmem16a* in E15 trachea examined by FISH. *Shh* marks tracheal epithelial cells. Nuclei are stained by DAPI (white). Scale bar indicates 20 μm. (B) Expression of *Cftr* and *Ano1* in tracheal epithelial cells at different time points, with a cell type breakdown for E15 and E16. P1 and P4 are shown in Figure S3A. (C) Cellular composition for tracheal epithelial cells of wild-type littermates and *Tmem16 a^-/-^* mutants from different time points. (D) Quantification of SCGB1A1^+^ secretory cells and Centrin-GFP^+^ ciliated cells of wild-type and *Tmem16a^-/-^* mutant trachea samples at P3. N=5 for each genotype. P-value (unpaired t-test) are indicated. Error bars represent standard deviation (S.D.). (E) Representative TEM images of secretory cells from wild-type and *Tmem16a^-/-^* mutant trachea at P0. Scale bar indicates 2 μm. Examples of large intracellular vesicles of the mutants are highlighted by asterisks. More examples are shown in Figure S3B. N=4 for each genotype. (F) Periodic acid-Schiff (PAS) Staining showing accumulation of mucus in wild-type and *Tmem16a^-/-^* mutant trachea at P3. The bronchial lumen is indicated by arrowheads in blue. Scale bar indicates 100 μm. (G) Differential interference contrast (DIC) images of flat-mounted trachea, flow path lines, and velocity magnitude of ciliary flow generated by wild-type and *Tmem16a^-/-^* mutant trachea samples at P2. Flow directions are indicated by arrows in red. Flow movie was taken at 1s/frame for 300 seconds. Flow movies played at 30 frame/second are included in Movie S1 and S2. X represents the medial-lateral axis, while Y represents the anterior-posterior axis. (H) Velocities of ciliary flow in wild-type and *Tmem16a^-/-^* mutant trachea samples at P2. N=3 for each genotype were included for analysis. p = 0.008 for |V|, p= 0.718 for V_x_, and p <0.001 for V_y_ (multiple t-test). Error bars represent S.D.

A handful of mouse models that lack *CFTR* have been generated to study the pathogenesis of cystic fibrosis, but they fail to develop CF features including mucus obstruction, mucus cell hyperplasia, as well as chronic and persistent airway infection and inflammation (Lavelle et al., 2016; McCarron et al., 2018). In contrast, genetic inactivation of *Tmem16a* results in tracheomalacia, similar to those observed in the CF animal models (Bonvin et al., 2008; Meyerholz et al., 2010; Rock et al., 2008), and shows accumulation of mucus in the trachea (Rock et al., 2009). We thus used *Tmem16a^-/-^* mice as an entry point to assess the role of chloride channels in airway development.

We processed mouse trachea of *Tmem16a KO* mutants and wild-type controls at E16, P1 and P4. We found an expansion of the secretory cell population in *Tmem16a^-/-^* mutants across all time points (Figure 3C). Immunofluorescent staining using postnatal trachea samples confirmed a significant increase in SCGB1A1^+^ cells in the mutants (Figure 3D). In contrast, a transgene reporter Centrin-GFP, which labels the basal bodies of motile cilia (Bangs et al., 2015; Higginbotham et al., 2004), indicated an insignificant reduction in the number of ciliated cells in the absence of *Tmem16a*.

Besides an increase in their population, secretory cells in *Tmem16a^-/-^* mutants suffer from alterations in their subcellular structure morphology. Specifically, secretory cells exhibited reduced microvilli, a dilated ER lumen and accumulation of large amorphous vesicles by TEM analysis (Figures 3E and S3B). Using Periodic acid–Schiff (PAS) and Jacalin staining, which stain mucosubstances such as glycoproteins, glycolipids and mucins (Matsuo et al., 1997; Ostedgaard et al., 2017), we observed strong signals in the trachea and bronchus of newborn *Tmem16a^-/-^* mutants, consistent with a mucus obstruction of the lower respiratory tract (Figures 3F and S3C). We next assessed tissue-level mucociliary clearance by characterizing the flow dynamics of fluorescent beads generated by airway motile cilia. At P2, wild-type trachea showed directional flow at 0.90 ± 0.22 μm/s from the distal to the proximal trachea. *Tmem16a* mutants showed minimal and sometimes reversed flow at a much lower speed of 0.42 ± 0.18 μm/s (Figures 3G and 3H; Movies S1 and S2). A combination of mucus cell hyperplasia and defects in ciliary clearance may account for mucus obstruction observed in the neonatal *Tmem16a^-/-^* mutants.

### *Tmem16a* KO Mutants Exhibit Abnormal Accumulation of *Krt4/Krt13* Secretory Cells

To probe into the cellular origin of mucus cell hyperplasia observed in *Tmem16a* mutants, we analyzed cell states of different epithelial populations from E16. At this stage, secretory cells include a basal-to-secretory transition state (BasalSecretory) and an immature secretory state (Secretort_Krt4), both of which are characterized by high expression of *Krt4*, a squamous cell marker (He et al., 2015) (Figures 4A and 4B). In addition, we found a more differentiated population expressing high levels of mature secretory markers, such as *Muc5b*, *Creb3l1* and *Gp2*, as well as enzymes involved in mucin glycosylation, such as *Galnt6* and *B3gnt6* (Figure 4B). Compared to wild-type mice, the percentage of cells from the Secretory_Krt4 state strongly increased in *Tmem16a* mutant mice. Given that *Krt13* is often paired with *Krt4* in epithelial cells (He et al., 2015), we confirmed the presence of *Krt4/Krt13* cells in mutant trachea via immunofluorescent staining of KRT13. Consistent with our transcriptomic analysis, KRT13 was barely visible in the postnatal control epithelium, but it showed strong signal in *Tmem16a* mutants (Figures S4A and S4B).

**Figure 4.**
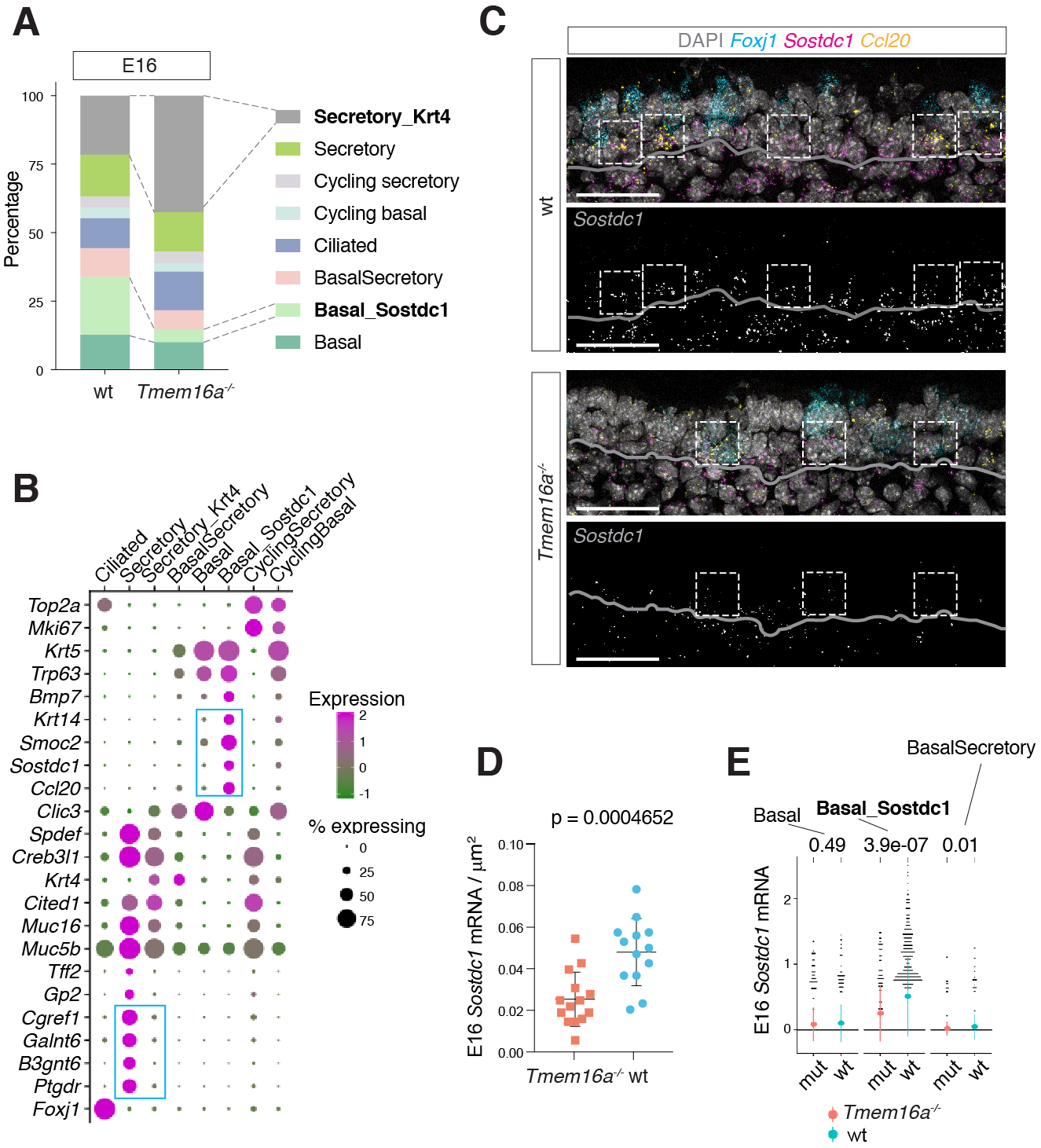
Altered Basal Cell Composition in *Tmem16a^-/-^* Mutant Trachea. (A) Cellular composition of tracheal epithelial cells from wild-type and *Tmem16a^-/-^* mutants at E16. (B) Dot plot depicting expression of marker gene of cell types shown in (A). Markers for Basal_Sostdc1 and secretory cells are highlighted in blue box. The size of the dot encodes the percentage of cells expressing the gene, while the color encodes the mean of expression level which has been normalized, log-transformed, and z-score transformed. Both wild-type and *Tmem16a^-/-^* mutants are included for this analysis. (C) Expression of *Sostdc1*, *Ccl20*, and *Foxj1* in wild-type and *Tmem16a^-/-^* mutant trachea samples at E16 detected by FISH. *Sostdc1* single channel images are shown in black and white. Grey lines indicate the basement membrane that separates epithelial cells from mesenchymal cells. Dashed squares indicate *Sostdc1^+^/Ccl20^+^* cells. Scale bar indicates 40 μm. (D) Concentration of *Sostdc1* RNA probes from wild-type and *Tmem16a^-/-^* mutant trachea samples at E16. N=3 for each genotype from each genotype were included in analysis. P-value (unpaired t-test) are indicated. Error bars represent S.D. (E) Expression level of *Sostdc1* in wild-type and *Tmem16a^-/-^* mutants at E16 in different populations of basal cells. Each dot represents a cell. Expression value has been normalized and log-transformed. Colored circles indicate mean expression values. Colored vertical lines cover the range of one standard deviation above or below the mean. Adjusted p-values for the comparison within each cell type (unpaired wilcoxon-test between wild-type and mutant) are indicated.

In the meantime, the proportion of tracheal basal cells of *Tmem16a* mutant mice is reciprocally decreased, suggesting a role of *Tmem16a* in the maintenance of the basal progenitor pool (Figures 4A). At E16, we identified a basal state expressing *Ccl20*, *Sostdc1*, *Smoc2*, and *Krt14* significantly higher compared to the rest of basal cells (Figures 4B and 4C). *Ccl20* belongs to the CC chemokine family and acts under both homeostatic and inflammatory conditions (Schutyser et al., 2003). *Sostdc1* is an antagonist of Wnt and Bmp signaling and plays a role in inhibiting the differentiation of epithelial progenitors (Närhi et al., 2012; Welsh and O’Brien, 2009). *Smoc2* is a calcium-binding protein expressed in *Lgr5*^+^ crypt progenitor cells (Muñoz et al., 2012). *Krt14* expression is high in airway basal progenitor cells required for tissue repair (Ghosh et al., 2013; 2011; Hong et al., 2004), and is upregulated in basal cell carcinoma (Rock et al., 2010). Taken together, this population of basal cells most likely represent a progenitor pool that gives rise to luminal cells during development and upon injury. In *Tmem16a* mutants, the percentage of *Sostdc1*^+^ basal cells was reduced at E16 (Figure 4A and 4C), and the expression level of *Sostdc1* was also significantly downregulated (Figures 4D and 4E), indicating a key role of *Tmem16a* in the basal-to-secretory cell fate transition.

### *Tmem16a* Does Not Affect the Developmental Trajectory of Ciliated Cells

In contrast to the paradigm of airway cell fate specification, in which Notch signaling regulates the balance between ciliated cells and secretory cells, we did not observe a significant change in the percentage of ciliated cells in the absence of *Tmem16a* (Figures 3C and 3D). To determine whether *Tmem16a* has any effect on the developmental dynamics of ciliated cells, we characterized the molecular composition of embryonic ciliated cells from both wild-type and *Tmem16a^-/-^* mutant tracheas.

At E16, ciliated cells consist of three distinct states that are all marked by *Foxj1*, including one (state I, Figure 5A) with high *Foxn4* expression, a transcriptional factor required for motile cilia formation in *Xenopus* epidermis and in cultured human bronchial epithelial cells (Campbell et al., 2016; Plasschaert et al., 2018). In addition to *Foxn4*, cells from state I also express *Ccno*, *Mcidas*, and *Gmnc*, indicating a precursor state of ciliated cells (Figure 5A). We next analyzed transcriptional trajectory of epithelial cells by assessing the ratio of unspliced to spliced reads via Velocyto, a method to estimate the rate of change of mRNA levels in time, aggregating over many genes to predict future transcriptional states for cells (La Manno et al., 2018). The trajectory of ciliated cells shown by the mRNA velocity model suggested that these *Foxn4*^+^ cells (I, Figures 5A and 5B) represented a nascent stage that would transition into more mature states expressing elevated *Cdhr3* (II and III, Figures 5A and 5B; Figure S5), consistent with the reported functional profiles of genes associated with each state. More mature states for ciliated cells involve a set of markers including *Sntn*, *Lrrc23*, a radial spoke protein required for vertebrate motile cilia function (Han et al., 2018), and *Prr18*, a proline-rich protein enriched in oligodendrocytes without further known insights into its function (Figure 5A).

**Figure 5.**
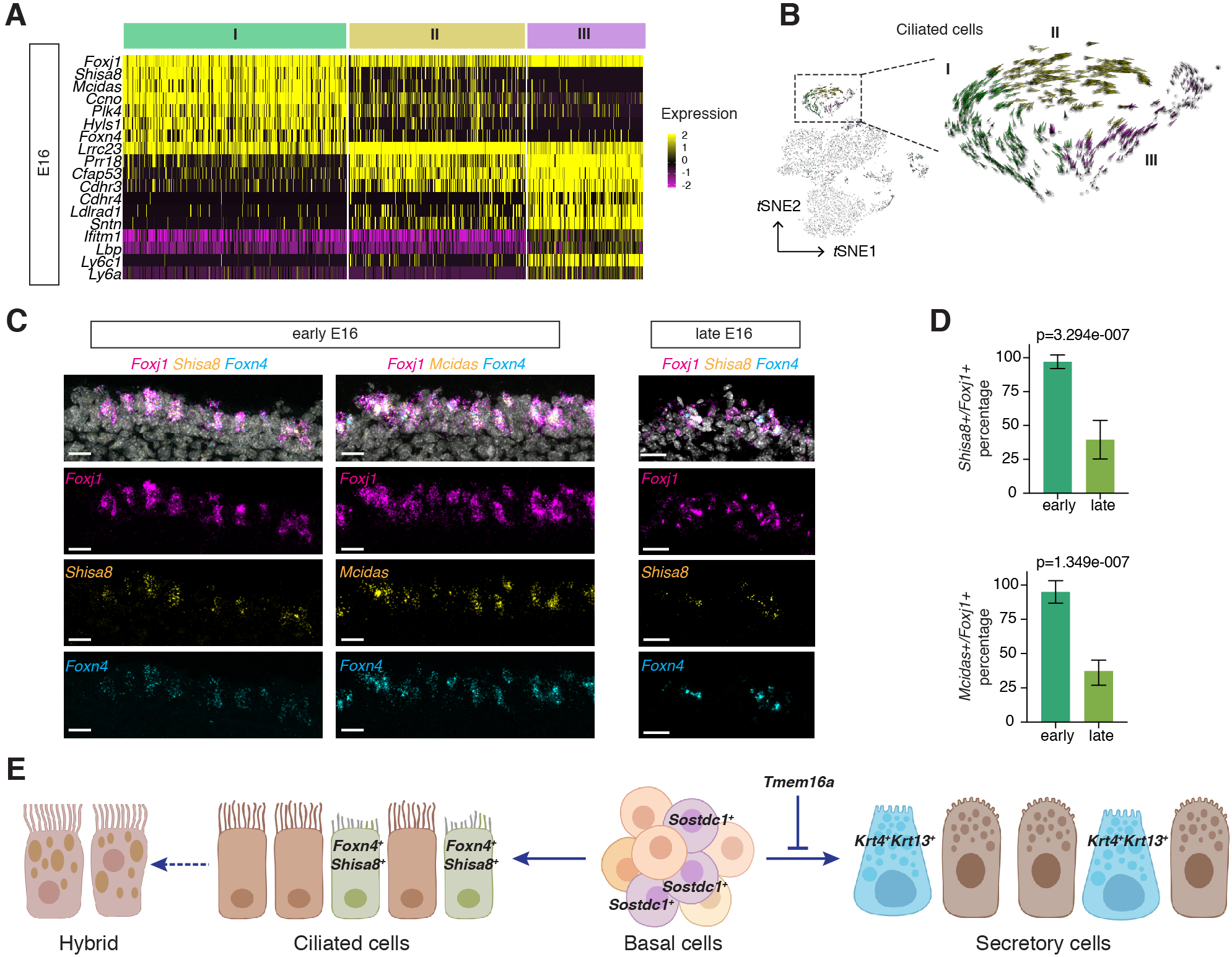
Developmental Dynamics of Ciliated Cells. (A) A heat map showing state-specific marker genes of ciliated cells at E16. Gene expression has been normalized, log-transformed, and z-score transformed. (B) Single cell velocity estimates for individual E16 tracheal epithelial cells. Arrows show the extrapolated states projected onto the *t*SNE plot for both wild-type and *Tmem16a^-/-^* mutant cells. mRNA velocity estimates for a selected list of genes are shown in Figure S5. (C) Expression of *Foxn4*, *Shisa8*, *Mcidas*, and *Foxj1* in early E16 and late E16 tracheal cells detected by FISH. Nuclei are stained by DAPI (white). Scale bar indicates 20 μm. (D) Quantification of *Shisa8*^+^ /*Foxj1*^+^ cells and *Mcidas*^+^ /*Foxj1*^+^ cells in early and late E16 trachea in the wild-type. Three littermates for each stage were included in analysis. Error bars represent S.D. (E) Model of epithelial cell differentiation of the embryonic trachea. *Tmem16a* is required to maintain a progenitor pool marked by *Sostdc1* and to inhibit secretory cell hyperplasia associated with an expansion of *Krt4*^+^/*Krt13*^+^ cells. The developmental trajectory of ciliated cells, including a progenitor state marked by *Foxn4* and *Shisa8*, are not affected by *Tmem16a* deficiency.

The precursor state of ciliated cells exhibits a novel and specific enrichment of *Shisa8*/*Ckamp39*, an auxiliary subunit for the AMPA receptor and a member of the CKAMP glutamate receptor family required for synaptic transmission (Farrow et al., 2015). FISH analysis of *Mcidas* and *Shisa8* showed colocalizations with *Foxn4* and *Foxj1* in E16 wild-type trachea (Figure 5C). The *Foxn4*-expressing precursor state appears to be under tight developmental control. In early E16 when dissections were performed early in the morning, all *Foxj1*^+^ cells were marked by *Foxn4*, *Shisa8*, and *Mcidas*. In late E16 when dissections were performed towards the end of the day, less than 40% of *Foxj1*^+^ cells were labeled by *Foxn4* and *Shisa8* (Figures 5C and 5D).

Overall, the percentage, transcriptome profiles, and developmental trajectory of ciliated cells are comparable between wild-type control and *Tmem16a* mutants. Based on the aforementioned changes in the cellular composition and molecular profiles between wild-type and *Tmem16a* mutant tracheal epithelial cells, we propose that *Tmem16a*-mediated chloride homeostasis maintains a *Sostdc1*-expressing progenitor pool within the epithelial basal cells, loss of which results in secretory hyperplasia at least partially due to an accumulation of intermediate secretory cells with elevated levels of *Krt4* and *Krt13* (Figure 5E). Lineage specification of ciliated cells, on the other hand, appears unaffected in the absence of *Tmem16a*.

### Immune Profiles of the Mucosal Barrier in Wild-type and *Tmem16a KO* Mutants

Epithelial cells of the respiratory tract are in direct contact with the external environment and form the front line of innate host defense by producing a diverse arsenal of antimicrobial molecules and cytokines (Ganesan et al., 2013; Vareille et al., 2011). Our analysis revealed the specific cellular source and temporal expression patterns of a set of immune modulators important for mucosal barrier function and airway immunity (Figure 6A). Based on their molecular profiles, these epithelial-derived molecules can be divided into two major groups. The first group consists of secreted peptides involved in the recognition of inhaled pathogens and mucin components (Figure 6A). For example, *Lbp* and *Ltf* encode secreted proteins that can bind and sequester lipopolysaccharides, a common cell wall component of Gram-negative bacteria and extremely strong stimulators of innate immunity (Actor et al., 2009; Bingle and Craven, 2004). *Defb1*, which encodes β-defensin, shows a broad spectrum of antimicrobial activity and acts as a multifunctional mediator in infection and inflammation (Ganz, 2003; Schneider et al., 2005). Mucin components, such as *Muc5b*, *Muc1*, and *Muc16*, as well as several surfactant proteins, such as *S*ftpa1 and *Sftpd*, provide lubrication and form a scaffold for the mucosal layer (Fahy and Dickey, 2010; Haczku, 2008). The second group includes cytokines and signaling molecules implicated in mucosal inflammatory responses (Figure 6A), such as *Cxcl15* (Chen et al., 2001), *Cxcl17* (Burkhardt et al., 2014), *Ccl28* (Hieshima et al., 2003), *Nfkbia* and *Nfkbiz*, which are inhibitors for the NF-κb pathway serving as a pivotal mediator of inflammatory responses (Liu et al., 2017; Zhang et al., 2017), as well as *Ptgs2*, *Ptges and Ptgds*, which are required for the biosynthesis of prostaglandin, a potent agent in the generation of the inflammatory response (Ricciotti and FitzGerald, 2011).

**Figure 6.**
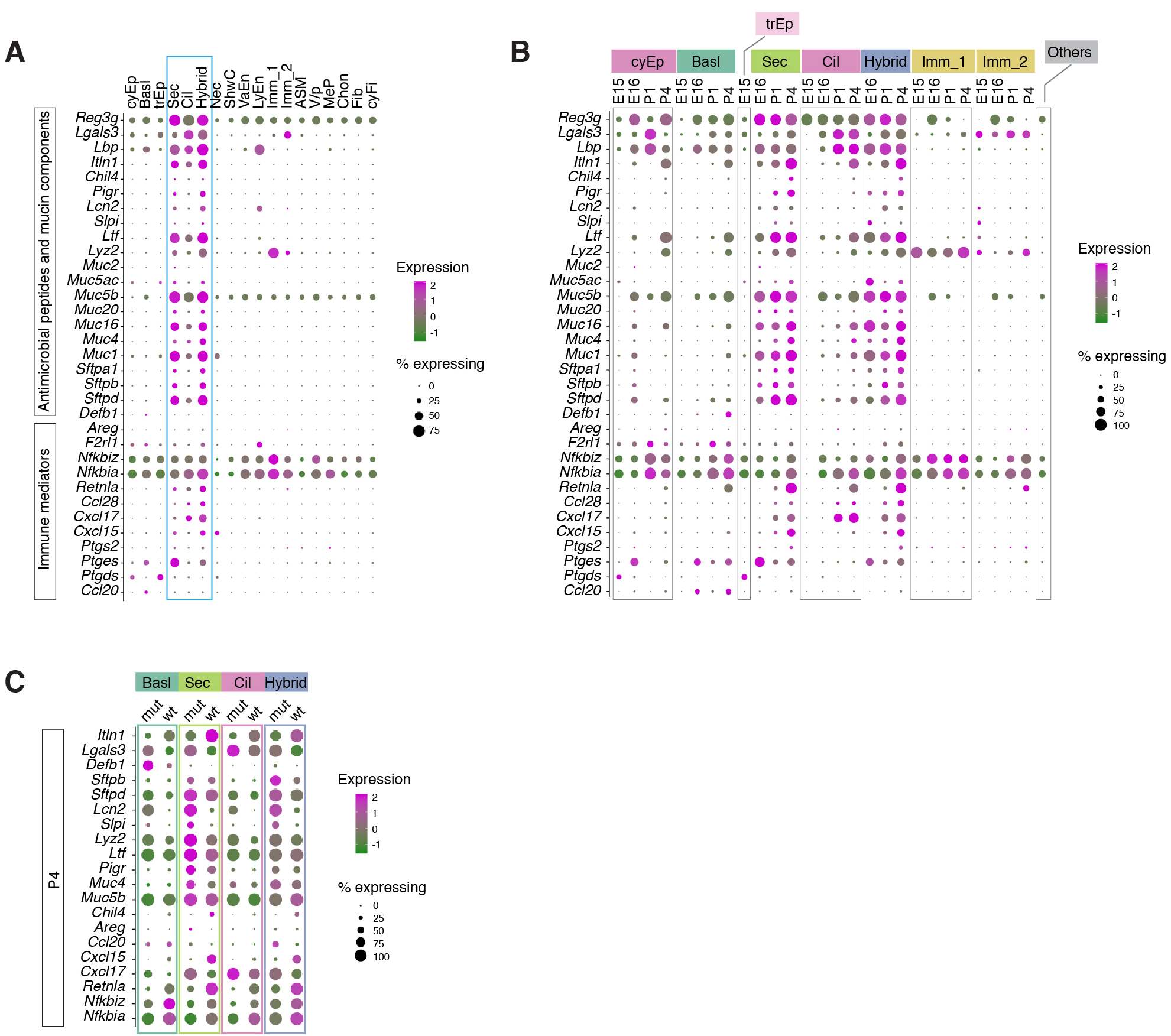
Immune Profiles of the Trachea Mucosal Barrier. (A) Dot plot showing the expression of mucosal barrier regulators and immune mediators. Secretory cells (Sec), Ciliated cells (Cil), and Cilia-Secretory Hybrid (Hybrid) collectively express high levels of these molecules and are highlighted in the blue box. The size of the dot encodes the percentage of cells expressing the gene, while the color encodes the mean of expression level which has been normalized, log-transformed, and z-score transformed. (B) Dot plot showing the expression of immune genes shown in (A) in different developmental time points. The size of the dot encodes the percentage of cells expressing the gene, while the color encodes the mean of expression level which has been normalized, log-transformed, and z-score transformed. (C) Expression of selected mucosal barrier regulators altered in *Tmem16a^-/-^* mutant epithelial cells at P4. The size of the dot encodes the percentage of cells expressing the gene, while the color encodes the mean of expression level which has been normalized, log-transformed, and z-score transformed. Expression profiles of the same genes at E16 are shown in Figure S6A.

Most, if not all, of these epithelial-derived molecules involved in the host defense and barrier function are produced by secretory and cilia-secretory hybrid cells. However, barrier function is not limited to these cells. For example, basal progenitor cells express *Defb1* while ciliated cells produce *Cxcl17*, *Lbp*, and *Lgals3*, which encodes galectin-3 and is implicated in pulmonary fibrosis in humans (Nishi et al., 2007) (Figure 6A and 6B). Both groups of immune cells express high levels of NF-κb inhibitors. Lymphatic vascular cells express *F2lr1*, known in humans as PAR2, and *Lcn2*, both of which are important modulators during homeostasis and inflammation (Joyal et al., 2014). Expression levels for many of these genes are age-dependent and peaked at P4 in our study, supporting the notion that the neonatal period is a critical period for establishing the airway mucosal barrier (Figure 6B).

In the neonatal epithelium, *Tmem16a* mutants exhibited a much reduced expression of immune regulators such as *Nfkbia/z*, *Retlna* and *Chil4* (Figures 6C and S6B). Both *Retlna* and *Chil4* are mediators of the Th2 inflammatory response (Nair et al., 2009; 2005; Pesce et al., 2009; Zhu et al., 2004). In contrast, antimicrobial molecules such as *Lgals3*, *Lcn2*, and *Ltf* are upregulated in *Tmem16a* mutants during embryogenesis, and the expression differences become more pronounced after birth (Figures 6C and S6B). The abnormal expression profiles of host-defense genes and immune modulators indicate that *Tmem16a* plays and important role for the formation of epithelial mucosal barrier.

### Mouse and Human Airways Share Similar Cellular Landscape during Development

To characterize cell types and gene modules during human airway development, we profiled ∼9700 cells from human fetal airway at gestation weeks 21 and 23 and (GW21 and GW23), establishing a single cell atlas for the human developing trachea (Figure 7A). While the tissue architecture of the human airway is more complex compared to mice, the core features of the airway landscape during embryogenesis are conserved between mice and humans (Figure 7B). From the human fetal samples, we identified orthologous cell types and states expressing similar sets of molecular programs as in our mouse atlas. For example, a population of mesenchymal cells expressing *CD34*, *THY1*, and *WNT2*, resembles the mesenchymal progenitors found in the mouse trachea. Other mesenchymal cell types, including vascular and lymphatic endothelial cells, chondrocytes, and airway smooth muscles, all share similar marker genes with their mouse counterparts (Figure 7B). In the epithelium, *FOXJ1*^+^ human ciliated cells exhibit a precursor state marked by *FOXN4* and *MCIDAS*, as well as a mature state marked by *SNTN* and *CDHR3*, consistent with our discoveries in mice (Figures 7A and S7A-B). Within basal cells marked by *TP63*, *SOSTDC1*, and *KRT14*, a subpopulation distinctive from the others expresses *SOX9* (Basal_SMG in Figure 7A), suggesting its origin from the submucosal gland (SMG) (Tata et al., 2018), a major structural difference between human and mouse trachea. The SOX9 negative basal population hence likely corresponds to apical basal cells which are located in the surface epithelium (SE) (Basal_SE in Figure 7B). Similarly, based on the expression of *SOX9*, *LTF*, *LRRC26*, *AQP5*, and *CCL28*, all of which are markers for the SMG (Ordovas-Montanes et al., 2018), we assigned the *MUC5B*-expressing secretory cells to the submucosal glands (Secretory_SMG in Figures 7A and 7B). The surface epithelial secretory cells were characterized by *SERPINB3* and *MUC16* that are typically found in surface epithelial cells (Secretory_SE in Figures 7A and 7B) (Fischer et al., 2009). Based on the expression of both the SMG marker *LTF* and the smooth muscle marker *ACTA2*, we annotated a cluster from the fetal trachea as myoepithelial cells, which are absent in mice.

**Figure 7.**
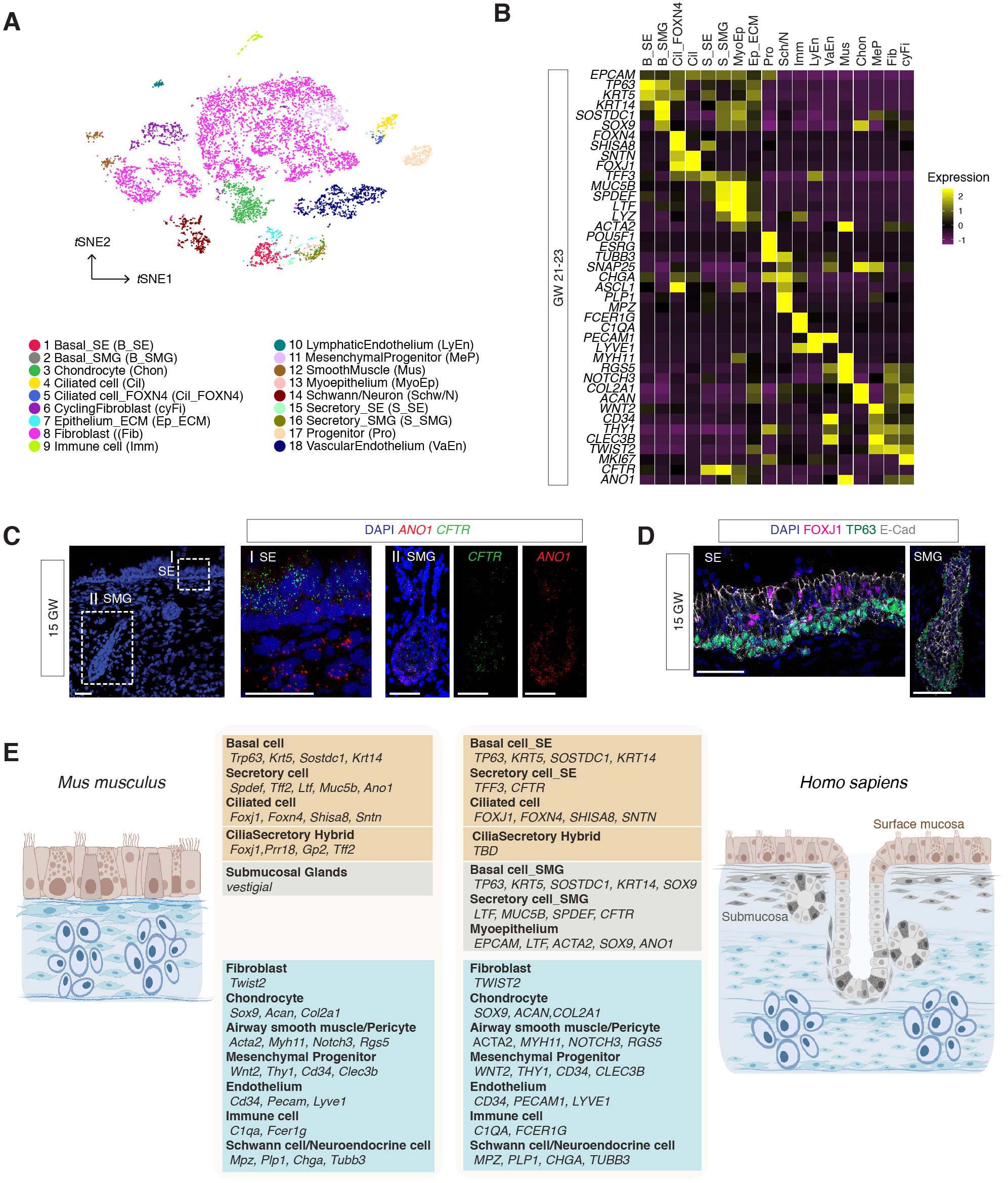
Cellular Composition and Molecular Profiles of Human Fetal Trachea. (A) Human fetal tracheal cells visualized on a *t*-distributed stochastic neighbor embedding (*t*SNE) plot. Colors indicate cell types and states identified in this study. (B) Heat map showing expression of marker genes for cell types identified in human fetal trachea of gestation week (GW) 21-23. Gene expression has been normalized, log-transformed, and z-score transformed. (C) Expression of *CFTR* and *ANO1/TMEM16A* in human fetal trachea of GW 15 by FISH. Two areas (I: surface epithelium/SE; II: submucosal glands/SMG) of the fetal tracheal epithelium are enlarged. Nuclei are stained by DAPI (blue). Scale bar indicates 50 μm. (D) Immunofluorescence staining of TP63 (green) and FOXJ1 (magenta) in human fetal tracheal sample at GW 15. SE and SMG are shown. Epithelial cells are marked by E-cad in white. Nuclei are stained by DAPI (blue). Scale bar indicates 50 μm. (E) Summary of cell types and marker genes reflecting similarity and distinction between mouse and human trachea.

Unlike being restricted to rare ionocytes in adults, *CFTR* mRNA was broadly distributed in the human fetal airway at GW15 (Figure 7C), consistent with the early manifestation of CF phenotypes seen in young patients. At this stage, epithelial basal cells start to differentiate based on the presence of *FOXJ1*^+^ cells (Figures 7D and S7B). The expression of *CFTR* becomes sparse at GW 21-GW 23, and is relatively restricted to secretory cells (Figure 7B). On the other hand, *ANO1/TMEM16A* transcripts are more abundant in myoepithelial cells (Figures 7B and 7C). The change in the expression pattern of *CFTR* during the course of human trachea development is reminiscent of the pattern of *Tmem16a* observed in the mouse developing airway, indicating that different chloride channels may play species-specific roles that are essential for airway development and mucosal barrier function. Together, our data identify *Tmem16a* KO mutant mice as a platform for studying chloride homeostasis in mammalian airway development and pathology.

## DISCUSSION

Cell fate specification and morphogenesis primarily take place during embryogenesis, and deviations from normal developmental processes can be detrimental to airway function in youth and may have long-lasting effects later in life. Here we present a comprehensive, high-quality single-cell atlas of the embryonic and neonatal mouse trachea, as well as the human fetal trachea, and systematically delineate the ontogeny of the large conducting airway and identify gene modules and cell types conserved between mice and humans. In the normal mouse developing trachea, we identified a rich repertoire of cell types, including epithelial cells, a large collection of mesenchymal cells, immune cells, endothelial cells, and neuronal cells that can be further divided into more fine-grained and novel cell states associated with distinct developmental stages and functional profiles, reflecting the structural and physiological complexity of the conducting airway. Importantly, all cell types identified in the mouse trachea have corresponding orthologous cell types in humans, indicating that the two species employ conserved gene regulatory programs to build the large airway during embryogenesis (Figure 7E).

Our assessment of enrichment patterns for selected airway disease genes using scGPS confirmed that most monogenic disease genes implicated in PCD are specifically expressed in ciliated cells. In contrast, susceptibility loci for COPD are expressed in both epithelial and stromal cell types, consistent with the pleiotropic presentations of complex-trait airway disease. The enrichment profiles generated by scGPS thus provide a framework for studying cell-type specific contributions in the pathogenesis of complex respiratory diseases.

Spanning four developmental time points, our dataset uncovers milestones of trachea development from the onset of differentiation, through cell fate determination during embryogenesis, and into the air breathing transition at birth. Compared to the homeostatic and regeneration phases in adults, the developing trachea exhibits both quantitative and qualitative differences in several key aspects of its transcriptional landscape. First, embryonic tracheal tissue expresses high levels of cell cycle markers and shows a higher percentage of progenitor cells, including the *Sostdc1*^+^/*Smoc2*^+^ basal progenitor cells located within the epithelium. Second, we identified gene modules for different cell states that may account for stage-dependent functional profiles. For example, a conserved precursor state for ciliated cells is marked by *Foxn4* and the novel marker *Shisa8*. *Foxn4* is required to promote motile cilia formation in *Xenopus* epidermis (Campbell et al., 2016), and its expression is very transient during frog embryogenesis (Briggs et al., 2018). Our data indicate that *Foxn4* expression peaks transiently at E16 in the mouse trachea as ciliated cells begin to emerge, supporting a central role of *Foxn4* in promoting motile ciliogenesis in mammals. Third, we uncovered two critical transcriptional events for the establishment of the mucosal barrier. During embryogenesis, epithelial cells upregulate multiciliated gene modules and secretory gene modules to initiate a massive differentiation process between E15 and E16. After the animals are born and start breathing, tracheal epithelial cells upregulate a set of mucosal cytokines, antibacterial effectors, and Th2 immune response genes that are critical for the maturation of barrier function.

Furthermore, we identified a cilia-secretory hybrid cell state that exhibits the combined molecular profiles of both ciliated and secretory cells. In the normal trachea, these hybrid cells start to emerge at E16 and become most abundant at P1. The hybrid state appears to be very rare during the adult homeostatic state and cannot be induced by chemically-triggered airway regeneration (Rawlins et al., 2007). In contrast, a cluster of cilia-secretory hybrid cells are found in the human inflammatory airway, indicating that neonatal airway may share similar immune status to the inflamed adult airway (Ordovas-Montanes et al., 2018; Vieira Braga et al., 2019); in both cases, hybrid cells are likely derived from a ciliated cell lineage. Supporting this notion, transdifferentiation of ciliated cells into goblet cells can be induced by IL-13, a major cytokine involved in allergy and inflammatory response (Turner et al., 2011; Tyner et al., 2006). While the *in vivo* functional attributes of this cell state have not been defined, it is possible that these hybrid cells reflect the airway plasticity and can fine tune the balance between efficient mucociliary clearance and mucus production.

Although the cell types identified in our study largely overlap with those identified in the mouse adult airway, we did not observe typical ionocytes that express both *Foxi1* and *Cftr* in this study. Instead, we showed that *Cftr* is ubiquitously expressed in trachea epithelial cells prior to differentiation and its expression is drastically decreased in postnatal airways. Similarly, *CFTR* is expressed in human fetal trachea and has been reported in other fetal mucosal epithelia, including the lung (Harris et al., 1991; McCray et al., 1992; Tizzano et al., 1993). *CFTR* in the adult airway is required for anion secretion in order to maintain osmotic balance for airway surface liquid (ASL) and mucus viscosity necessary for effective mucociliary clearance (Bertrand and Frizzell, 2003; Button et al., 2012; Cozens et al., 1994). Because the fetal airway is constantly exposed to amniotic fluid, it remains unclear whether *CFTR* during embryogenesis is required to facilitate fluid secretion analogous to its role in the adult airway.

Consistent with the early expression of *CFTR*, congenital airway defects have been reported in CF fetuses, raising the possibility that chloride homeostasis may play a role in airway development (Gosden and Gosden, 1984; Larson and Cohen, 2005; Ornoy et al., 1987). In this study, we took advantage of a *Tmem16a* KO mouse model as an entry point to test the possible involvement of chloride channels in airway development. We show that *Tmem16a* in mice and *CFTR* in humans share similar temporal and spatial expression within the developing tracheal epithelium, and that genetic inactivation of *Tmem16a* in mice recapitulates hallmarks of CF symptoms. Through a combination of transcriptomic analyses and *in vivo* characterization, we identified two key mechanisms of *Tmem16a* underlying murine airway development and barrier functions, respectively. First, *Tmem16a* in the undifferentiated epithelial cells plays an essential role in maintaining the progenitor pool of tracheal basal cells by limiting the differentiation of basal cells into the secretory lineage. Such activity of *Tmem16a* may be sufficient to account for the severe mucus cell hyperplasia observed in *Tmem16a^-/-^* mutants. Second, persistent expression of *Tmem16a* in differentiated airway epithelial cells controls the normal status of the neonatal mucosal immunity. A shift in the expression of antimicrobial genes and proinflammatory modules in *Tmem16a* mutants indicates that these mutants may be more prone to infection and inflammation. Intracellular chloride, regulated by chloride channels, has been implicated in vesicle trafficking and plays a role in regulating plasma membrane dynamics (He et al., 2017; Stauber and Jentsch, 2013). While the specific actions of *Tmem16a* in basal cell differentiation and barrier functions require further investigation, it is possible that *Tmem16a* mediated-chloride homeostasis may underlie both aspects of *Tmem16a* required for airway development and physiology.

Mouse embryonic and human fetal trachea share common cell types and genetic markers (Figure 7E). Inactivation of *CFTR* in humans and *Tmem16a* in mice, lead to congenital abnormalities of the airway, raising the possibility that perhaps intracellular chloride homeostasis, modulated by these two chloride channels, represents a conserved component for mammalian airway development. Although the pathogenesis and clinical presentations of CF in humans are extraordinarily complex, and the mucosal immunities differ among species, our study provides a conceptual foundation for the ontology of mammalian trachea, and demonstrates the presence of conserved cell types and gene modules present in the mouse and human conducting airway. Notwithstanding the challenge in reconciling the discrepancies in mouse and human CF pathogenesis, our tractable mouse model allows for discoveries of airway cell types that require chloride channels for proper differentiation and functions that are relevant to early onset airway diseases.

## Acknowledgements

We thank James Webber from Chan Zuckerberg Biohub for support with RNA-seq data processing; Laurence Baskin of UCSF for providing access to human fetal samples; Kathryn Anderson of Sloan Kettering Institute for sharing Arl13bmCerry/Centrin-GFP transgenic mice; the Bios Core Histology and Biomarker Teams at UCSF for histology analysis; Tong Cheng, Guillermina Ramirez-san Juan, and Jorge Alexis Vargas for reagents and tissue analysis. This work is supported by Chan Zuckerberg Biohub and the National Institutes of Health (R01NS069229 to L.Y.J. and F32HD089639 to M.H.). L.Y.J. and Y.N.J. are investigators of the Howard Hughes Medical Institute.

## Author Contributions

Conceptualization, M.H and B.W

Methodology, M.H and B.W

Software, B.W. and D.L.

Validation, M.H and B.W

Formal Analysis, M.H., B.W., D.L., W. Y. and Y.C.

Investigation, M.H., B.W., W. Y., V.P., Y.C., K.L., R.S and M.T

Resources, A.S., N.N. and M.C

Data Curation, B.W. and S.D

Writing – Original Draft, M.H.; Writing – Review and Editing, M.H., B.W., Y.C., S.D. and L.J

Visualization, M.H and B.W

Funding Acquisition, M.H., N.N., Y.J., S.D. and L.J

## Methods

### Mice

The *Tmem16a* null allele, *Ano1^tm1Jrr^*, has been described previously (Rock et al., 2008). Breeding colonies were maintained in a mixed genetic background by outcrossing C57BL/6J *Ano1^tm1Jrr^* males to FVB females, which were obtained from JAX. For lineage tracing, *Shh^tm1(EGFP/cre)Cjt/J^* ((Harfe et al., 2004) and *Gt(ROSA)26Sor^tm4(ACTB-tdTomato-EGFP)Luo/J^* (Muzumdar et al., 2007) were obtained from JAX. Arl13b-mCheery/Centrin-GFP reporter mice *Tg(CAG-Arl13b/mCherry)^1Kv^*and *Tg(CAG-EGFP/CETN2)^3-4Jgg/KvandJ^* (Bangs et al., 2015) were obtained from Sloan Kettering Institute. Mice were housed in an animal facility and maintained in a temperature-controlled and light-controlled environment with an alternating 12-hour light/dark cycle. A maximum of five mice were housed per cage. All protocols have been approved by the University of California San Francisco Institutional Animal Care and Use Committee.

### Isolation of mouse trachea cells

To obtain embryonic tracheal cells for scRNA-seq, pregnant female mice were sacrificed via CO_2_ asphyxia at desired stages and embryos were collected. Neonatal mice were sacrificed by decapitation. Tracheas were collected and washed with ice cold DMEM/F12 1:1 (Gibco, 25200056) to remove residual blood. For single cell dissociation of embryonic trachea, samples were dissociated with 0.25% Trypsin-EGTA and 0.1 mg/mL DNaseI in DMEM/F12 at 37 °C for 15 minutes. For neonatal trachea, samples were incubated with a combination of 1 mg/mL elastase (Worthington, LS006363) and 2.5 mg/mL dispase II (Roche, 4942078001) in DMEM/F12 0.1 mg/mL DNaseI for 15 minutes, and then 0.125% Trypsin-EGTA for 15 minutes at 37 °C. Digest reaction was terminated by an addition of 10% bovine calf serum. Dissociated single cell solution was centrifuged at 300g for 5 minutes at 4°C. Cell pellets were resuspended with cold DMEM/F12 with 5% bovine calf serum and passed through a 35 µm filter into a collection tube. Single cells were counted with a Neubauer chamber and cell viability was assessed with trypan blue staining. Using this protocol, we consistently obtained > 90% viable cells. For cells expressing GFP and RFP dissociated from *mT/mG* samples were separated collected via Fluorescence Activated Cell Sorting (FACS) using SH800S (Sony) sorters. Post-sorting, the cells were collected in cold DMEM/F12 media with 5% FBS till library preparation.

### Isolation of human fetal tracheal cells

Human fetal trachea samples were obtained and used in accordance with the guidelines for the care and use of animals and human subjects at University of California, San Francisco. Given that deidentified fetal tissue were involved, this study does not involve human subjects as defined by the federal regulations summarized in 45 CFR 46.102(f) and does not require IRB oversight. Details for approval is included in the accompanying GDS certification letter from the UCSF Ethics and Compliance and the Human Research Protection Program with study ID number 16-19909. First and early second trimester human fetal trachea were collected without patient identifiers after elective termination of pregnancy with approval from the Committee on Human Research at UCSF (IRB#12–08813). Fetal age was estimated using heel-toe length (Drey et al., 2005). Fetal age was calculated from time of fertilization, fetal age, and not from last menstrual period. Fetal trachea samples were collected at room temperature during surgery and analysis (1-2 hours) before stored in at 4 degrees C. For dissociation, tissue pieces were rinsed in ice cold PBS and incubated in a combination of 200 μg/ml Liberase (Liberase™ TL Research Grade, Roche, 05401020001) and 0.1 mg/mL DNaseI in DMEM/F12 30 minutes. Digest reaction was terminated by addition of 10% bovine calf serum, followed by an additional step to remove red blood cells via RBC Lysis buffer (ThermoFisher). Following steps were identical to the one used for mouse single cell collection.

### Single cell RNA sequencing

Single cells were encapsulated into emulsion droplets using the Chromium Controller (10x Genomics). scRNA-seq libraries were constructed using Chromium Single Cell 3’ reagent kits v2 (mouse samples) or v3 (human samples) according to the manufacturer’s protocol. About 3000 to 7000 cells were targeted and loaded in each channel. Reverse transcription and library preparation were performed on C1000 Touch Thermal cycler with 96-Deep Well Reaction Module (Bio-Rad). Amplified cDNA and final libraries were evaluated on Agilent tapestation system (Agilent Technologies). Libraries were sequenced with 100 cycle run kits with 26 (v2) or 28 (v3) bases for Read1, 8 bases for Index1, and 98 bases (v2) or 91 bases (v3) for Read2 on the Novaseq 6000 Sequencing System (Illumina) to over 80% saturation level.

### Antibodies and immunostaining

Antibodies for immunofluorescence staining were mouse anti-FOXJ1 (1:500, 2A5, ThermoFisher, 14-9965-82), rabbit anti-TRP63/P63 (1:500, proteintech, 12143-1-AP), SCGB1A1/CC10 (1:200, B6, Santa Cruz, sc-390313), rabbit anti-KRT13 (1:500, proteintech, 10164-2-AP), mouse anti-acetylated α-tubulin (1:2,000; 6–11B-1; Sigma-Aldrich T6793), rat anti–E-cadherin (1:1,000; ECCD-2; Thermo Fisher Scientific), rabbit anti-DCLK1 (1:500, ThermoFisher, PA5-20908), Alexa Fluor 488-, 594- and 633-conjugated secondary antibodies (Invitrogen), and Fluorescein labeled Jacalin (1:500, Vector Laboratories, FL-1151). For protein immunostaining, cells or tissue sections were fixed with 4% paraformaldehyde (PFA) for 20 min at room temperature or −20C° methanol for 10 minutes on ice. After fixation, the samples were washed and blocked with IF buffer (1× PBS with 1% heat-inactivated goat/donkey serum and 0.3% Triton X-100). Primary antibodies were added and incubated for 1 h at room temperature or overnight at 4 °C. After washing with IF buffer, secondary antibodies and DAPI were added at 1:1,000 dilution for 1 h at room temperature. Samples were washed with 1× PBS and mounted with Fluoromount-G (SouthernBiotech). Washing and staining were performed with IF buffer (1× PBS with 5% serum and 0.2% triton) at room temperature. Samples were then imaged using a Leica TCS SP8 confocal microscope with the 40× and 63× HC PL Apo oil CS2 objective.

### Transmission electron microscopy

For transmission electron microscope, embryonic and newborn tracheas were dissected in cold PBS and fixed with 2% paraformaldehyde and 2.5% glutaraldehyde in 0.1 M sodium cacodylate buffer. After buffer rinses, samples were postfixed in 1% OsO_4_ at room temperature for 4 h followed by dehydrating in an ethanol series. Samples were stained with osmium tetroxide and embedded for thin sectioning in EPON. Sections of 70–100 nm were examined on a JEOL transmission electron microscope and photographed at primary magnifications of 4,000–30,000X.

### Histology

Newborn trachea and lung were fixed by formalin, dehydrated, and embedded in paraffin. Standard Periodic acid–Schiff (PAS) staining for airway mucus was formed at Mouse Pathology Core at the University of California, San Francisco.

### Cilia flow analysis

For imaging of ciliary flows, tracheas were dissected in ice-cold Dulbecco’s Modified Eagle Medium: Nutrient Mixture F-12 (ThermoFisher, DMEM/F12), and sliced open along the proximal-distal axis prior to imaging. Each trachea was mounted onto 35mm cell imaging dishes (MatTek) with a solution of fluorescent beads (Carboxylate-modified Microspheres 0.4 μm size; Invitrogen, 1:250 to 1:500) in 500ml DMEM/F12 media. Imaging was performed in an environmental chamber with 5% CO_2_ at 37C° and acquired using a Leica TCS SP8 confocal microscope with a10x HCX PL Apo dry CS objective. Flow was imaged at 1f/s for 300s. Flow pathlines were generated using Flowtrace (Gilpin et al., 2017) (http://www.wgilpin.com/flowtrace_docs/). Particle Image Velocimetry (PIV) fields were generated using PIVLab (https://www.mathworks.com/matlabcentral/fileexchange/27659-pivlab-particle-image-velocimetry-piv-tool) for MATLAB. We used the FFT window deformation PIV algorithm with three passes consisting of 128×128, 64×64, 32×32 interrogation areas to determine the velocity vectors. Velocity vectors were filtered around a Gaussian Distribution (within 0.8 standard deviation) around no movement to remove incorrect pairings and extraneous movements. All parameters were held constant across all analyses to reduce systematic errors due to inconsistent discretization.

### mRNAs fluorescent *in situ* hybridization (FISH)

*In situ* hybridization for mouse *Ano1*, *Shh*, *Cftr*, *Foxj*1, *Foxn4*, *Gp2*, *Prr1*8, *Shisa8*, *Mcidas1*, *Sostdc1*, *Ccl20*, and human *ANO1*, *CFTR*, *FOXJ1*, and *FOXN4* were performed using the RNAscope kit (Advanced Cell Diagnostics) according to the manufacturer’s instructions.

### Statistical analysis

Methods for statistical analysis and numbers of samples measured in this study are specified in the figure legends. The error bars indicate the SD.

### Data analysis

Sequences generated by the NovaSeq were de-multiplexed and aligned to the mm10.1.2.0 genome using CellRanger (10x Genomics) with default parameters. Subsequent filtering, variable gene selection, reduction of dimensionality, clustering, and differential expression analysis with Wilcoxon rank sum tests were performed using the Seurat package (version 2.3) in R.

### scGPS (single-cell Geneset percentile scoring)

The geneset percentile score for each cell was calculated as the mean of cell-wise percent rank for all genes in a certain module. For instance, a given module consists of the number of m genes, an 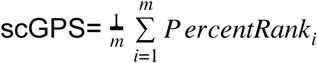, in which *PercentRank_i_* is the rank of the log-normalized *i*=1 expression level of *gene_i_* in this cell compared to *gene_i_* expression in all cells in the dataset. Equal values of *gene_i_* expression are assigned the lowest rank. Each ranking is scaled to [0,1]. An scGPS of p can be interpreted as the mean expression of the selected genes is p percentile for the given cell. Implementation in R can be found in (https://github.com/czbiohub/BingWu_DarmanisGroup_TracheaDevTmem16a).

### Velocyto analysis of ciliated cell dynamics

RNA velocity was estimated by following velocyto.py documentation. Spliced and unspliced transcript counts were derived from Cellranger’s outputs and with “run10x” default settings through velocyto.py command-line interface. Cell type annotations were predetermined as described in Figure 4 and Figure 5 with Seurat in R. Scripts to reproduce the results for E16 epithelial cell dynamics including the transition trajectory of ciliated cells are available at (https://github.com/czbiohub/BingWu_DarmanisGroup_TracheaDevTmem16a).

### Data and code availability

Sequencing reads and processed data in the format of gene-cell count tables are available from the Sequence Read Archive (SRA). All code used for analysis in this study is available on GitHub (https://github.com/czbiohub/BingWu_DarmanisGroup_TracheaDevTmem16a).

### Illustration

All cartoon illustrations were created with Biorender.

---

**Figure S1.**
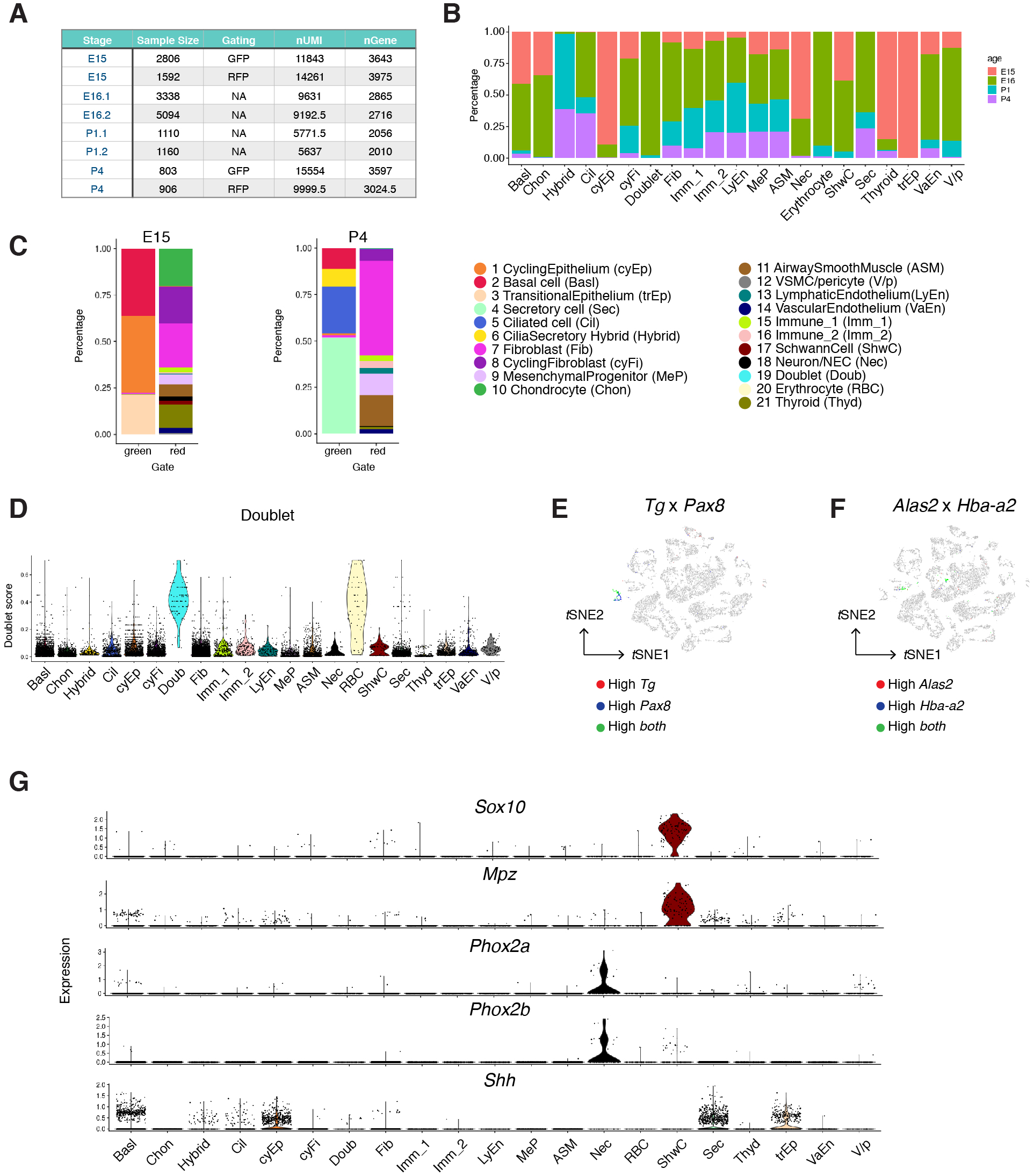

**Figure S2.**
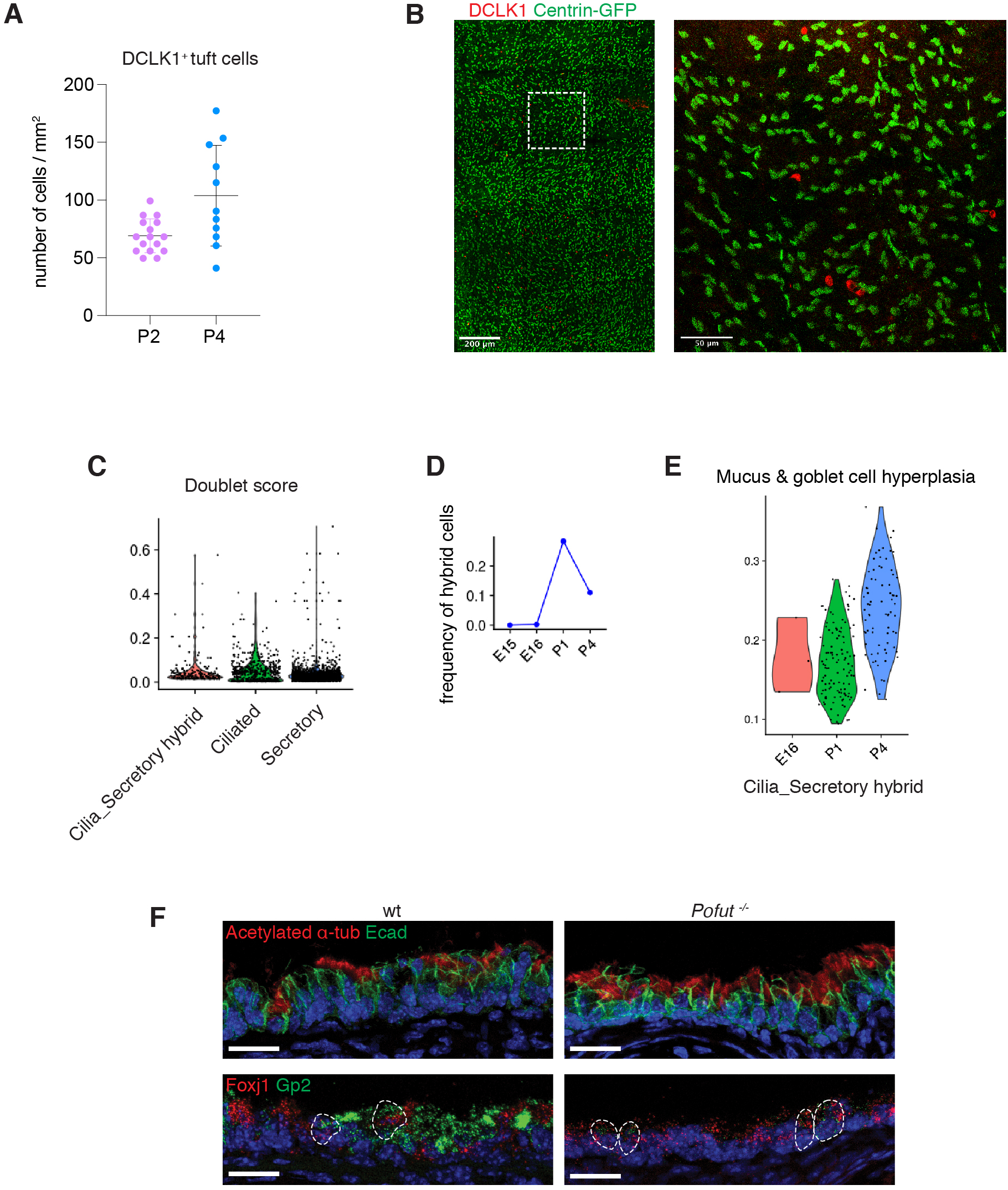

**Figure S3.**
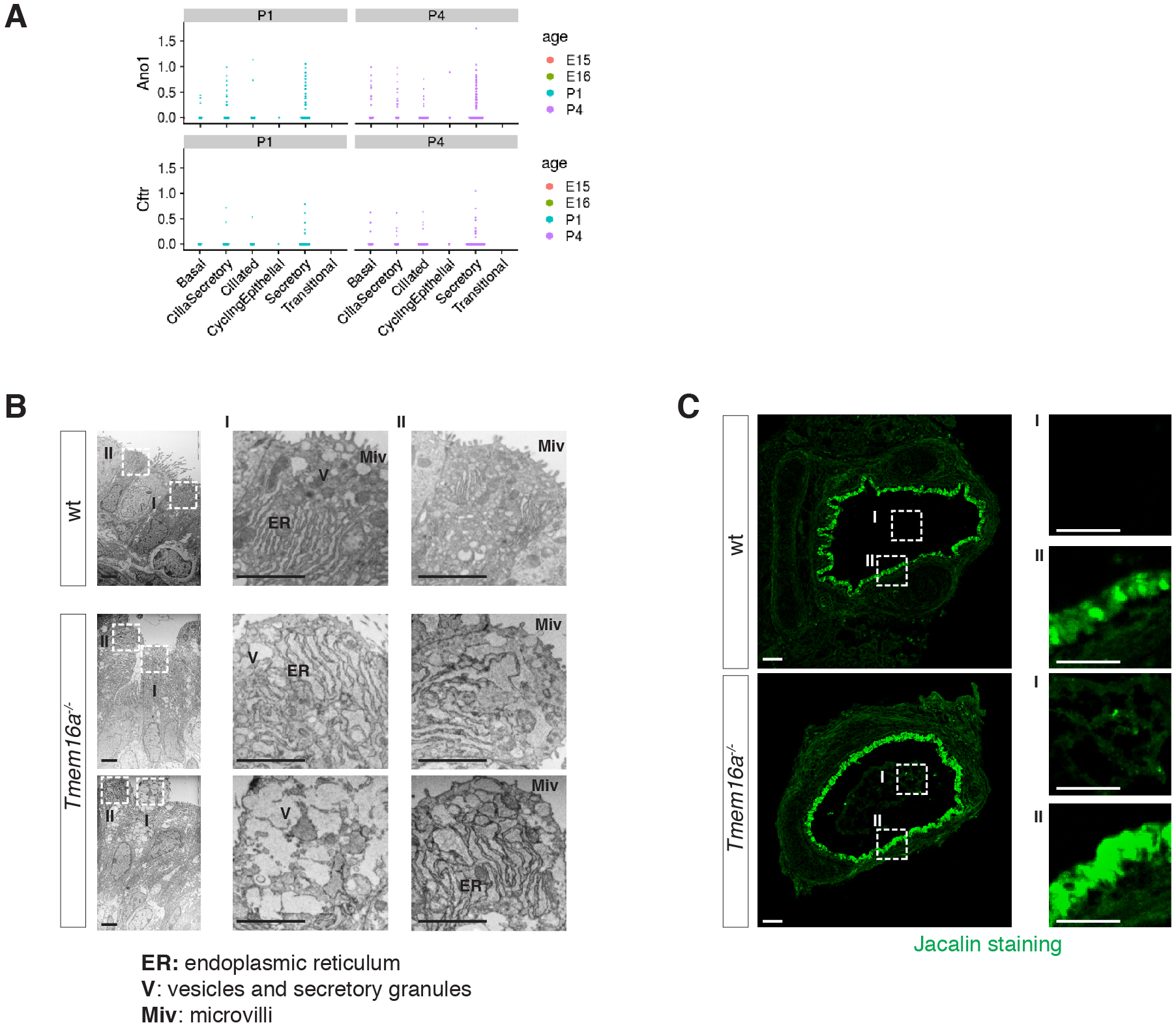

**Figure S4.**
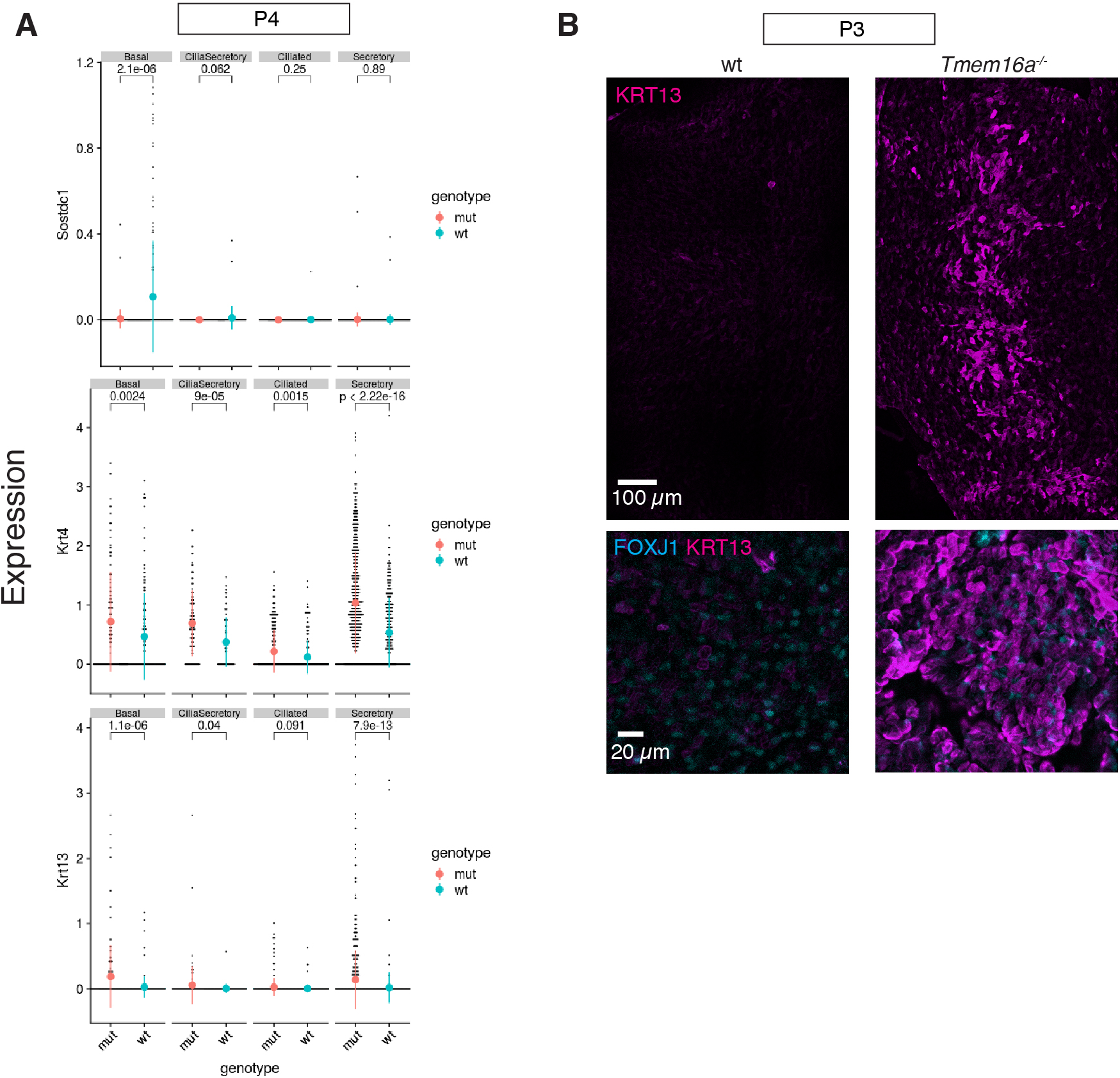

**Figure S5.**
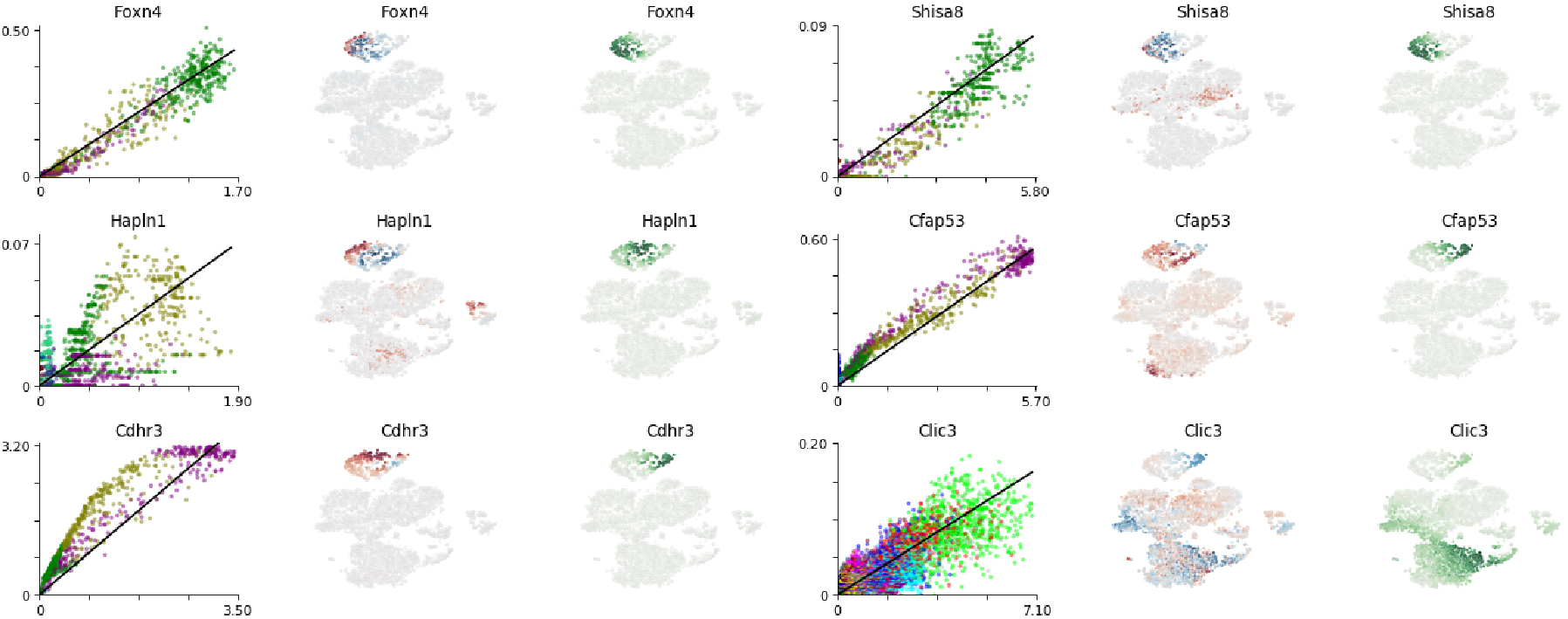

**Figure S6.**
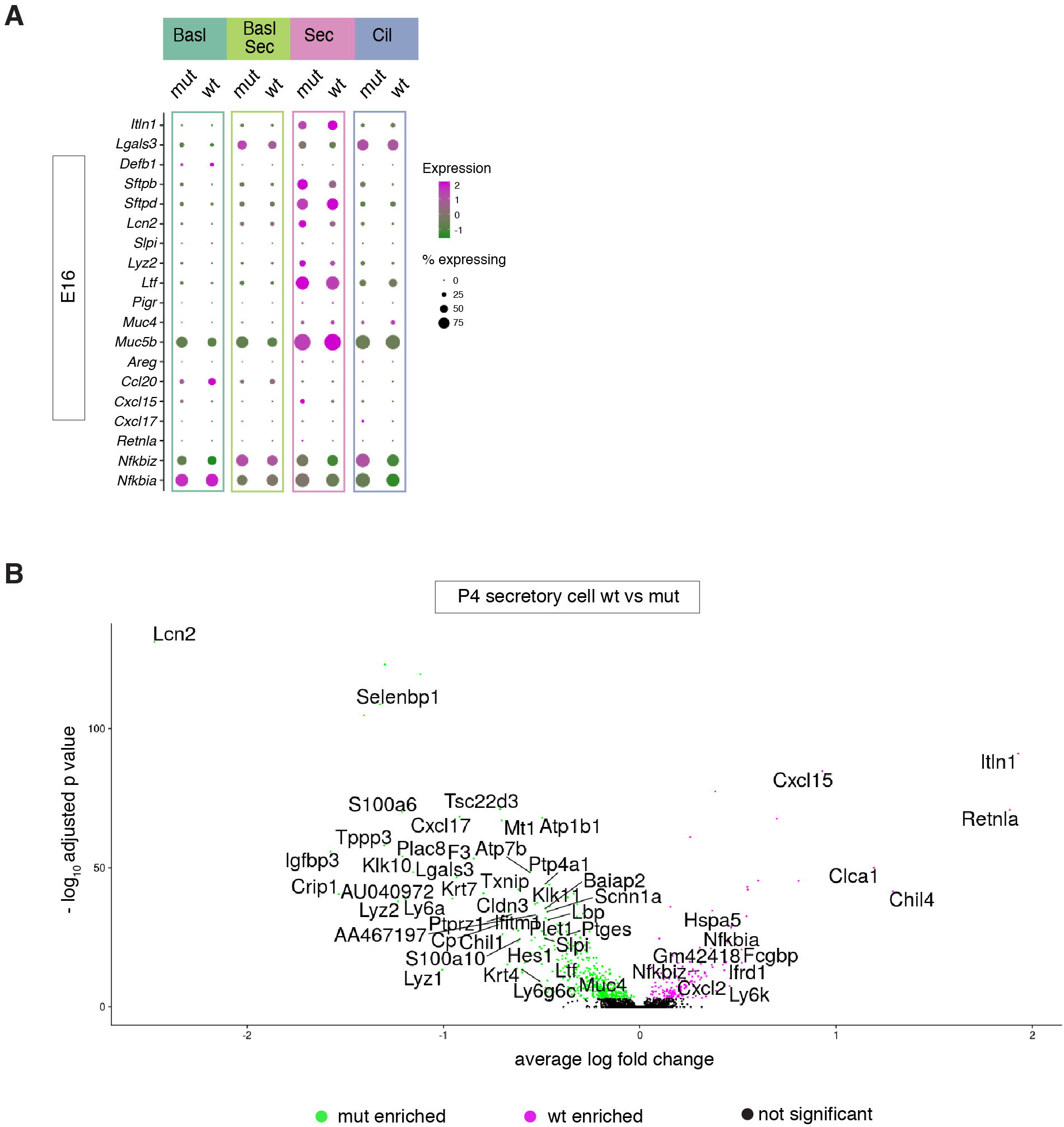

**Figure S7.**
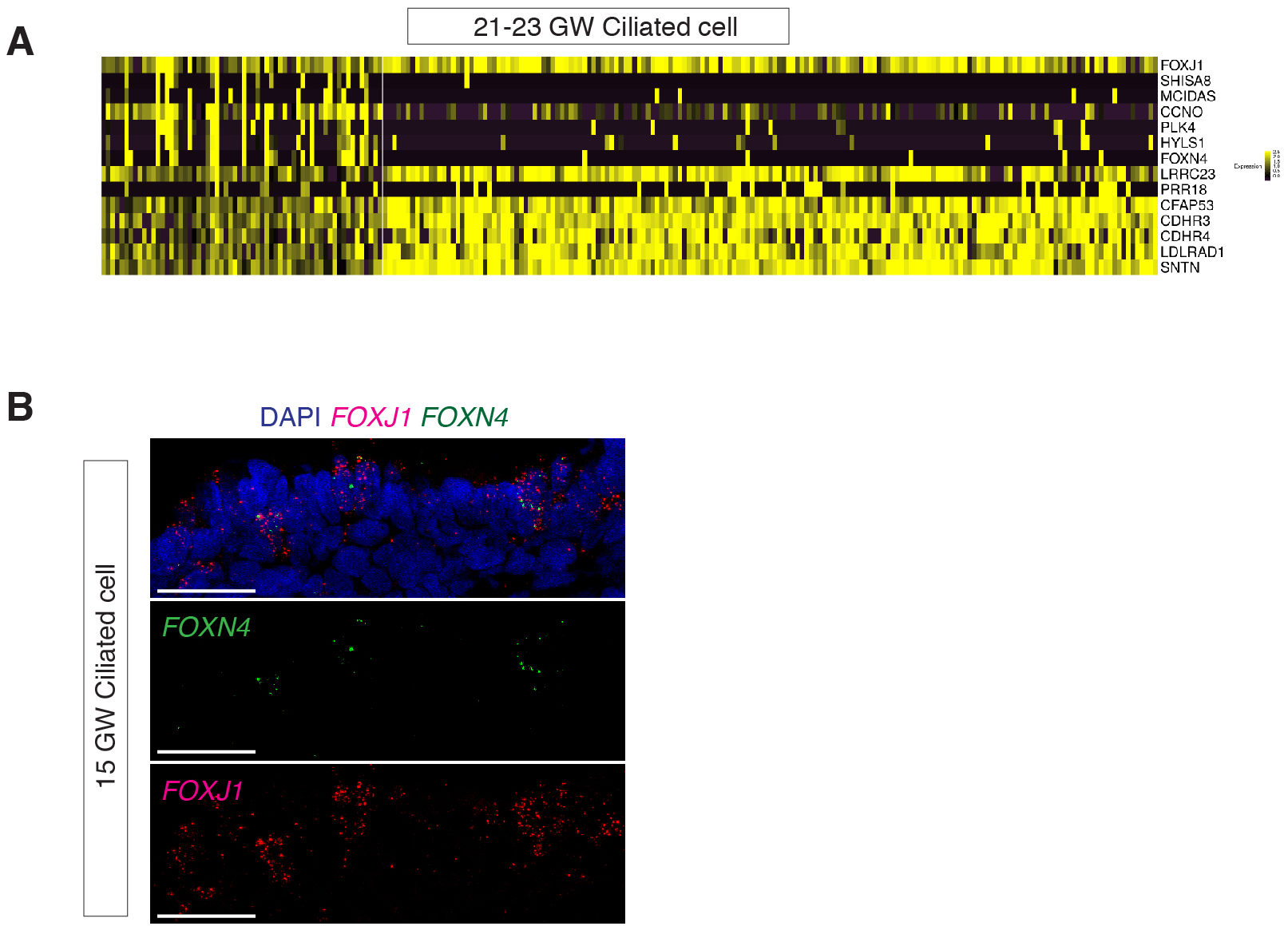

