## Supplement figure legends for "Single Cell RNAseq Reveals A Critical Role of Chloride Channels in Airway Development"

**Figure S1-7 Legends**

**Figure S1 Construction of the Developing Mouse Trachea Atlas, Related to Figure 1**
(A) A summary of cell numbers, gating of fluorescence-activated cell sorting (FACS, performed for E15 and P4 samples), median number of unique molecular identifiers (nUMI), and median number of genes (nGene) for each sample in the wild-type mouse trachea atlas presented in Figure 1B.
(B) Vertical bar graphs showing the proportion of cells from each time point for each tracheal cell type and states colored by developmental stages.
(C) The cellular composition of all cell types identified at E15 and P4. FACS was performed for cells from $Shh^{Cre}/R26^{mT/mG}$ mice before RNA-sequencing to distinguish cells of the Shh-expressing endoderm lineage (green) from the rest (red).
(D) Doublet scores for all cells. Each dot represents a cell. Colors indicate cell clusters.
(E) Expressions of thyroid markers *Tg* and *Pax8* projected onto *tSNE* shown in Figure 1B.
(F) Expressions of erythrocyte markers *Alas2* and *Hba-a2* projected onto *t*SNE shown in Figure 1B.
(G) Violin plots showing expressions of lineage markers for Schwann cells (Schw), Neuroendocrine cells (Nec), and endoderm cells in all cell types and states annotated. Gene expression has been normalized and log-transformed.

**Figure S2 Characterization of Tuft Cells and Cilia-Secretory Hybrid Cells, Related to Figure 2**.
(A) Numbers of tuft cells, marked by DCLK1, of P2 and P4 wild-type trachea. N= 3 from each stage were included for analysis. Error bars represent S.D.
(B) Distributions of ciliated cells, marked by Centrin-GFP, and Tuft cells, marked by DCLK1, in P4 wild-type trachea. *en face* view of the tracheal epithelial surface is shown. The numbers of tuft cells are orders of magnitude lower than the numbers of ciliated cells. Scale bars are indicated.
(C) Doublet scores for ciliated, secretory, and hybrid cells.
(D) Fraction of hybrid cells within the airway epithelium at E15, E16, P1 and P4. Representation of epithelial cell populations are shown in Figure 2A.
(E) scGPS for genes implicated in mucosal goblet cell hyperplasia for cilia-secretory hybrid cells at E16, P1, and P4. The gene list is included in Table S1.

(F) Hybrid cells in the trachea of *Pofut1*$^{-/-}$ mutant and wild-type littermates at P3. Compared to littermate controls, abundant ciliated cells with motile cilia, marked by acetylated-$\alpha$ tubulin, were present in *Pofut1*$^{-/-}$ mutants. *Foxj1* and *Gp2* double positive cells, highlighted by dashed circle, were present in both genotypes. Note that *Pofut1*$^{-/-}$ mutants lack cells that are only *Gp2*$^{+}$ . Scale bar indicates 20 mm.

**Figure S3 Involvement of *Ano1/Tmem16a* in Mucus Cell Hyperplasia of the Mouse Trachea, Related to Figure 3**.

(A) Expression of *Cftr* and *Ano1* in tracheal epithelial cells at different time points, with a cell type breakdown for P1 and P4. Expression of *Cftr* and *Ano1* at E15 and E16 are shown in Figure 3A.

(B) TEM images of wild-type and *Tmem16a*$^{-/-}$ mutant tracheal epithelial cells. Mutant secretory cells show reduced microvilli (Miv) and abnormal intracellular organizations, including dilated ER lumen (ER) and accumulation of vesicles (V). Scale bars indicate 2.5 $\mu$m.

(C) Jacalin-Alexa488 staining of wild-type and *Tmem16a*$^{-/-}$ mutant trachea at P0 labels secretory cells and glycoproteins. Inset I shows tracheal lumen; mucosubstances labeled by Jacalin-Alexa488 were apparent in the mutant trachea lumen. Inset II shows secretory cells. Scale bar indicates 50 $\mu$m.

**Figure S4 Expansion of *Krt4*$^{+}$/*Krt13*$^{+}$ Secretory Cells in *Tmem16a*$^{-/-}$ Neonatal Trachea, Related to Figure 4.**

(A) Expression levels of *Krt4*, *Krt13*, and *Sostdc1* in epithelial cell types in wild-type and *Tmem16a*$^{-/-}$ trachea at P4. Each dot represents a cell. Expression value has been normalized and log-transformed. Colored circles indicate mean expression values. Colored vertical lines cover the range of one standard deviation above or below the mean. Adjusted p-values for the comparison within each cell type (unpaired wilcoxon-test between wt and mutant) are indicated.

(B) Immunofluorescent staining of KRT13 and FOXJ1 in wild-type and *Tmem16a*$^{-/-}$ trachea at P3. Scale bars are indicated.

**Figure S5 Single-cell Velocity Estimates for Individual Ciliated Cells at E16, Related to Figure 5.**

Selected phase portraits and fits of the equilibrium slope for the epithelial cells at E16. For each gene, the first column shows spliced-unspliced phase portrait with the equilibrium slope fit shown by the line. The second column represents the magnitude of the residuals. The third column shows the expression for the spliced molecules.

**Figure S6 Altered Immune Profiles in *Tmem16a*$^{-/-}$ Mutant Epithelial Cells, Related to Figure 6.**

(A) Expression of selected mucosal barrier regulators altered in *Tmem16a*$^{-/-}$ mutant epithelial cells at E16. The size of the dot encodes the percentage of cells expressing the gene, while the color encodes the mean of expression level which has been normalized, log-transformed, and z-score transformed. Expression profiles of the same genes at P4 are shown in Figure 6C.
(B) Differential expression and associated significance for each gene (dot) between wild-type and *Tmem16a*$^{-/-}$ mutant secretory cells at P4. Green dots indicate *Tmem16a*$^{-/-}$ mutant enriched genes, magenta dots indicate wild-type enriched genes. Genes with a p-value above 0.001 are in black (not significant).

**Figure S7 Precursor State of Ciliated Cells Found in Human Fetal Trachea, Related to Figure 7.**

(A) A heat map showing state-specific marker genes of ciliated cells at human fetal stage 21-23 GW. Gene expression has been normalized, log-transformed, and z-score transformed. (B) FISH validation of *FOXN4* expression, co-localized with a subset of *FOXJ1*$^{+}$ cells, in human fetal trachea at 15 GW. Scale bars indicate 20 $\mu$m.

**Table S1 Gene Lists for scGPS analysis for airway diseases associated genes, Related to Figures 1E.**

**Table S2 Gene Lists for scGPS analysis for cell cycle scoring, Related to Figure 2C, and full list of genes used to generate heat maps of epithelial cells, Related to Figure 2D.**

**Movie S1 Flow Movie made from wild-type trachea.**

Ciliary flow generated by luminal ciliated cells across the entire trachea. Flow was imaged by Leica SP8 confocal microscopy at 1frame/s for 300s. Movie is shown at 30f/s. Fluorescent beads serve as tracer particles.

**Movie S2 Flow Movie made from *Tmem16a*$^{-/-}$ mutant trachea.**

Ciliary flow generated by luminal ciliated cells across the entire trachea. Flow was imaged by Leica SP8 confocal microscopy at 1frame/s for 300s. Movie is shown at 30f/s. Fluorescent beads serve as tracer particles.
